# A targetable developmental program co-regulates angiogenesis and immune evasion

**DOI:** 10.1101/2024.12.21.628442

**Authors:** Pietro Berico, Amanda Flores Yanke, Fatemeh Van Rajabpour, Catherine Do, Irving Simonin Wilmer, Theodore Sakellaropoulos, Ines Delclaux, Robert Stagnitta, Estefania Vázquez-Cruz, Iman Osman, Jane Skok, Carla Daniela Robles-Espinoza, Amanda W. Lund, Markus Schober, Eva Hernando

## Abstract

Ultraviolet (UV)-induced DNA mutations produce genetic drivers of cutaneous melanoma initiation and numerous neoantigens that can trigger anti-tumor immune responses in the host. Consequently, melanoma cells must rapidly evolve to evade immune detection by simultaneously modulating cell-autonomous epigenetic mechanisms and tumor-microenvironment interactions. Angiogenesis has been implicated in this process; although an increase of vasculature initiates the immune response in normal tissue, solid tumors manage to somehow enhance blood flow while preventing immune cell infiltration. By comparing the expression of transcription factors (TFs) across early-stage melanoma, naevi, and other cancer types, we found the homeodomain-containing TF HOXD13 drives a melanoblast-like developmental program, which is upregulated in melanoma and strongly correlated with angiogenesis and immune cell exclusion. Using transcriptomics, 3D chromatin profiling, and in vivo models, we demonstrate that HOXD13 upregulation promotes tumor growth in vivo by concomitantly enhancing angiogenesis and suppressing T-cell infiltration. HOXD13 orchestrates 3D chromatin contacts between distal enhancers and promoters, simultaneously activating VEGFA, SEMA3A, and CD73. VEGFA and SEMA3A remodel the tumor vasculature and CD73 elevates extracellular levels of adenosine, a vasodilator and immune suppressor that binds adenosine receptors (AdR) on endothelial and T cells. In line with these findings, HOXD13-induced growth advantage in vivo was significantly reversed by the concomitant administration of VEGFR and AdR inhibitors. By revealing a dual pro-angiogenic and immunosuppressive HOXD13-CD73/VEGF gene regulatory axis, we identify a subset of patients who might benefit from combinations of AdR and VEGFR inhibitors which are both currently being tested in clinical trials.

## INTRODUCTION

Efficient physiological immune responses require an increase in tissue vasculature. Paradoxically, many solid tumors find ways to simultaneously activate angiogenesis to support their increasing demands for oxygen and nutrients^1^while excluding immune cells^2^. The tumor vasculature’s morphology, maturation, and activity have recently emerged as significant determinants of immune cell homing, activity, and immunotherapy response^3–5^. However, the molecular mechanisms by which cancer cells govern an immune-refractory angiogenic process are largely unknown. In this context, cutaneous melanoma (SKCM) represents an exquisite study model. Due to the immunogenicity of neoantigens caused by the accumulation of UV-dependent mutations^6–9^, melanoma cells are challenged by high immune surveillance and a poorly vascularized epidermal-dermal junction beginning already at the earliest phases of tumorigenesis. Therefore, melanoma cells must rapidly invade the underlying dermis to escape from immune responses and access blood vessels that fuel their growth. Although the progression from a radial to vertical melanoma growth stage has been associated with angiogenesis and lymph angiogenesis^10^, its role in progression remains controversial because tumor vascular density does not predict patient outcomes^11^ . Meanwhile, tumor infiltrating lymphocytes (TILs) are strong predictors of patient outcomes^12^ , but they are not currently included in the American Joint Committee on Cancer (AJCC) melanoma staging system. Although the oncogenic hyperactivation of mitogen-activated protein kinase (MAPK) pathway, the most common SKCM tumorigenic driver, promotes angiogenesis and suppresses the immune response^13, 14^, it is not possible to predict which patients will recur and/or metastasize based on their somatic mutation profiles.

Here we explore whether transcriptional processes could play a role in regulating angiogenesis and immune evasion to drive melanoma progression and metastasis. By analyzing the expression of >1000 transcription factors (TFs) in primary melanomas and naevi^15^, we discover a highly significant association between increased Homeobox-containing TF HOXD13 expression, angiogenesis and immune cold melanomas. HOXD13 expression is not restricted to specific phenotypic melanoma cell states, suggesting that its activation precedes the emergence of intra-tumor heterogeneity, or its regulation is independent of cell state-defining transcriptional programs. Our loss-of-function and gain-of-function experiments in both human and syngeneic mouse models show that HOXD13 binds highly interconnected distal enhancers to simultaneously activate the expression of modulators of angiogenesis (*SEMA3A*, *VEGFA*^16, 17^) and the immune response (*NT5E). The NT5E* gene encodes the CD73 protein, a 5’-ectonucleotidase known to increase extracellular levels of adenosine, which simultaneously regulates the permeability of the vasculature^18, 19^ and suppresses immune cells^20^ by binding adenosine receptors (AdRs) on those cells. Accordingly, we show that HOXD13 enhances tumor growth by modulating both vessel maturation and immune cells infiltration. Moreover, the concomitant inhibition of angiogenesis and adenosine pathways synergistically abrogates the tumor growth advantage provided by HOXD13 by increasing immune infiltration. Our study exposes a developmental gene expression program that melanoma cells activate irrespective of their transcriptional cell state (i.e., melanocytic vs mesenchymal) to improve their fitness by concurrently modulating their vascular and tumor immune microenvironments. Our results could be impactful as they provide a rationale for treating immune cold HOXD13-positive melanomas by concomitantly blocking CD73 and VEGF/SEMA3.

## MATERIALS AND METHODS

### Cell culture

Primary melanocytes NHEM1, NHEM2 and a-HEM were purchased from PromoCell and cultured in Ham-F10 supplemented with 10% Fetal Bovine Serum (FBS), 2mM L-Glutamine, 200nM TPA, 200pM Cholera Toxin, 10ng/ml hSCF, 10nM EDN-1 and Penicillin Streptomycin (PS). Other cell lines (i.e., A375, B16F10, 293T, HCmel1274, YUMM1.7, MeWo) were purchased from the American Type Culture Collection (ATCC); 501mel from Yale University; SKMEL-5, SKMEL-28, SKMEL-239 and SKMEL-147 were kindly provided by Alan Houghton (Memorial Sloan-Kettering Cancer Center, New York, NY, USA); WM115, WM3211 were a gift from Meenhard Herlyn (Wistar Institute, Philadelphia, PA, USA); 113/6-4L cells (hereafter 4L, a derivative of WM239A) were provided by Robert S Kerbel and William Cruz-Munoz (Sunnybrook Research Institute, Toronto, Canada). Short-term cultures (STC) 12-273LN and 12-273BM were generated by Dr. Iman Osman^21^; Jurkat cells were kindly provided by Dr. Iannis Aifantis (NYU Grossman School of Medicine, Perlmutter Cancer Center, New York, NY, USA); D4M-3A were obtained from Dr David Mullins (Geisel School of Medicine, Dartmouth, NH, USA). 501mel, A375, MeWo, B16F10, 293T and 4L were cultured in high glucose DMEM supplemented with 10%FBS and PS. SKMEL cells were cultured in EMEM supplemented with 10%FBS and PS. WM cells were cultured as previously described^22^. YUMM1.7 and D4M-3A were grown in Ham-F12/DMEM supplemented with 10%FBS, 1% non-essential amino acids (NEAA) and PS. HCmel1274 and Jurkat were cultured in RPMI supplemented with 10%FBS, 2mM L-Glutamine and PS. All cell lines were grown at 37°C with 5% CO_2_ except for melanocytes which were cultured in 10% CO_2_.

All cell lines were tested negative for mycoplasma and monitored using MycoAlert® PLUS Mycoplasma Detection Kit (Lonza, LT07-703).

### Cell transfection and lentiviral infection

A375, SKMEL-5 and B16F10 cells were infected with custom made lentivector LT3GEPIR (Addgene, 111177) carrying doxycycline-inducible shRNAs against human or murine *HOXD13*, mRNAs following published protocols^23^. A375, YUMM1.7 and D4M-3A cells were infected with lentiviral vector pCW57-MCS1-2A-MCS2 (Addgene, 71782) EMPTY or engineered to express GFP or 3xFLAG-3xHA-hHOXD13 (GenScript). B16F10^OVA^ and HCmel1274^OVA^ were generated using the lentivector pLVX-puro-cOVA (Addgene, 135073). Jurkat^NY-ESO^ were generated by infecting a custom-made lentivector obtained by cloning a synthetized NY-ESO-TCR-P2A-GFP cDNA (Genscript) into an empty LV-EF1a-PuroR vector provided by Dr. Irwin Davidson (IGBMC, Department of Functional Genomics and Cancer, Illkirch, France).

For A375 and Jurkat co-culture, A375^GFP^ and A375^HOXD13^ were additionally infected with lentivector LV-RFP (Addgene, 26001). Engineered cell lines were selected using 3ug/ml of puromycin selective agent or sorted by flow cytometry. In vitro doxycycline induction experiments were conducted using 1ug/ml of doxycycline for at least 48hrs.

Transfection was carried out using Lipofectamine RNAiMAX (Thermo Fisher Scientific, 13778150) following manufacture instructions and using an siRNA final concentration of 25nM.

### Lentivirus production

293T cells (ATCC) were co-transfected with lentivector, psPAX2 (Addgene, 12260) and pMD2.G (Addgene, 12259) in Fugene6 (Promega, E2691). After 3 days of incubation, 293T supernatants were harvested, filtered and the virus concentrated 100 times 20k rpm for 2hr at 4°C using a Beckman Coulter Optima Ultracentrifuge. Then, concentrated virus was added to cells supernatant serum-free in presence of 5-10ug/ml of Polybrene (Sigma Aldrich, TR-1003-G) for 24hr. Afterward, cells were expanded for 3 days in complete media before selection in puromycin or FACS sorting.

### T cell isolation from OT-I mice

Eight to ten weeks old C57BL/6-Tg (TcraTcrb)1100Mjb/J or OT-I (The Jackson Laboratory, 003831) mice were humanely euthanized and the axillary, inguinal, brachial, mediastinal lymph nodes and the spleen were harvested in PBS. Afterward, lymph nodes and spleen were smashed with a syringe plunger on top of a 70uM strainer and 50ml conical tube and, after PBS rinsing, processed tissues were pelleted at 500g for 10min at 4°C. Next, red blood cells were removed by incubating the cell pellet with ACK buffer (Thermo Fisher Scientific, A1049201) for 1min at room temperature (RT). Final pellet was cultured and activated for 1 day in RPMI + 10%FBS + 50nM 2-Mercaptoethanol + 2mM L-Glutamine + 1ng/mL OVApeptide 257-264 (Genscript, RP10611) + PS. T cells were expanded in RPMI + 10%FBS + 50nM 2-Mercaptoethanol + 2mM L-Glutamine + 100U/ml hIL-2 (STEMCELL Technologies, 78036) + PS and kept in culture for 5-8 passages.

### Murine and human melanoma cells co-culture with T cells

For B16F10^OVA^ and HCmel1274^OVA^, 10k cells were transfected with either siCTRL or siHoxd13 in 96-well plates and after 72hr (timepoint in which cells reach ∼50k), 500k OT-I activated T cells were stained with CellTrace Violet (Thermo Fisher Scientific, C34557) and add to melanoma cells for 24hrs prior FACS analysis for AnnexinV+.

For A375^RFP^ co-culture with Jurkat^NY-ESO^, 5k melanoma cells were transfected with either siCTRL or siHoxd13 and seeded in 96well plates + IFN-γ (20ng/ml) for 48hrs (timepoint in which cells reach ∼50k number) prior addition of 500k Jurkat cells for 72hrs. For A375^RFP^ EMPTY and HOXD13, cells were induced with doxycycline during the whole experiment incubation. Cell density was measured using Incucyte (Sartorius) RFP+ signal.

### Proliferation assay

Cell growth assays have been conducted by seeding between 5-10k cells per well in 96-well plates and incubated for 72-96hrs in Incucyte. Each condition was represented by at least 3 technical replicates. Relative cell density was calculated based on the mean cell surface occupancy at the first time point of measurement (every 2-12-24hrs).

### ELISA assay

ELISA quantification of adenosine and VEGFA in the phenol red-free culture media was performed using Adenosine assay kit (Abcam, ab211094) and VEGF Quantikine ELISA kit reaction (R&D Systems, DVE00) respectively, and following the manufacturer’s instructions. Briefly, after 48hrs of doxycycline treatment or siRNA transfection, culture media was harvested and filtered with 0.45μm syringe filter (Corning, 431220) before ELISA quantification. DMEM phenol red-free culture media (Gibco, 21063-029) was used to prevent autofluorescence.

### Flow cytometry

For apoptosis quantification, cells were co-stained with AnnexinV-FITC (BD, 556420) and Propidium iodide (BD, 556463) in FACS binding solution (10mM Hepes pH7.4, 140mM NaCl, 2.5mM CaCl_2_) for 15min at RT in the dark. For CD73 protein surface analysis, cells were stained with anti-CD73 human (BioLegend, 127212) or mouse (BioLegend, 344012) in FACS buffer (1x PBS, 0.5% BSA, 5% FBS) for 30min at RT in the dark. Apoptotic cells (AnnexinV+), necrotic cells (AnnexinV+, PI+) and CD73 positive cells were measured using a BD FACS LSRII.

(Ghost Dye TMRed 780, Tonbo Biosciences; #13-0865-T500) followed by Fc receptor blockade with Fc Shield (Tonbo Biosciences; #70-0161-U500). For surface markers, single-cell suspensions were incubated with primary antibodies diluted in PBS containing 1% bovine serum albumin (BSA) for 30–45 min at 4 °C. Samples were run through BD FACSymphony A5 and data were acquired using BD FACSDiva software. Analyses were performed using BD FlowJo software (v10.10).

### RT-qPCR

Total RNA was purified using Nucleospin RNA Plus (Macherey-Nagel, 790498.250) following manufacture instructions. Next, between 50ng-1ug of purified RNA was converted to cDNA using Super Script IV synthesis kit (Thermo Fisher Scientific, 18090200) following manufacture instructions. Afterward, cDNA was diluted 10-300 times in water and qPCR was performed using PowerSYBR Green PCR Master Mix and measured using a Biorad CFX384 machine. mRNA relative quantification was calculated using comparative Ct method (Λ1Λ1Ct).

### Western Blot

Whole cell extracts and protein fractionations were performed as previously described^24^. For SDS-PAGE experiments, between 10-20ug of protein lysate were loaded with 4x Laemmli buffer (100mM Tris pH8, 0.4DTT, 4% SDS, 0.2% bromophenol blue, 30% glycerol) on pre-casted Nupage 4-12% gradient gels (Thermo Fisher Scientific, NP0322BOX, NP0323BOX). Proteins were transferred into nitrocellulose membrane (Cytiva, 10600011) and primary antibodies stainings were conducted overnight at 4°C. Secondary antibodies were incubated 1hr at RT and followed by alkaline phosphatase exposure using an Odyssey machine.

### Immunofluorescence and Immunohistochemistry

Mice tumors were placed in biopsy cassettes (Fisherbrand, 15182702E) and fixed for 72hrs in 10% Formalin fixative solution (EKI-Chem, 2406420) with gentle steering at room temperature. Then, tumors were rinsed and stored in 70% ethanol at 4°C before paraffin embedding.

Next, tissues were dehydrated through escalating grades of ethanol and xylene and infiltrated with paraffin (Paraplast X-tra, Leica SKU 39603002) on a Leica Peloris II tissue processor. Embedded tissues were sectioned at 5 µm and stained with hematoxylin (Leica, 3801575) and eosin (Leica, 3801619) on a Leica ST5020 automated stainer to evaluate histology. Adjacent sections were immunostained on a Leica BondRx autostainer, according to the manufacturers’ instructions. In brief, sections were first treated with peroxide to inhibit endogenous peroxidases followed by an antigen retrieval step with either ER1 (Leica AR9961) or ER2 (Leica AR9640) at 100o. For standard immunohistochemistry, sections were incubated with an antibody against the HA-tag (CST, cat # 3724S; clone C29F4; RRID 1549585) at a 1:200 dilution for 60 minutes at room temperature. Primary rabbit antibodies were detected with anti-rabbit HRP-conjugated polymer and 3,3’-diaminobenzidine (DAB) substrate that are provided in the Leica BOND Polymer Refine Detection System (Cat # DS9800). For multiplex immunofluorescence staining with Akoya Biosciences® Opal™ reagents, slides were incubated with the first primary antibody (see supplementary reagents) and secondary polymer (Rabbit-on-Rodent HRP polymer; Biocare RMR622), and then underwent HRP-mediated tyramide signal amplification with a specific Opal® fluorophore. The primary and secondary antibodies were subsequently removed with a heat retrieval step, leaving the Opal fluorophore covalently linked to the antigen. This sequence was repeated with subsequent primary and secondary antibody pairs and a different Opal fluorophore at each step (see table below for reagent details). Sections were counterstained with spectral DAPI (Akoya Biosciences, FP1490) and mounted with ProLong Gold Antifade (ThermoFisher Scientific, P36935). Semi-automated image acquisition was performed on either a Hamamatsu Nanozoomer whole slide bright field scanner at 40X magnification or on a Akoya Vectra Polaris (PhenoImagerHT) multispectral imaging system at 20X magnification using PhenoImagerHT 2.0 software in conjunction with Phenochart 2.0 and InForm 3.0 to generate unmixed whole slide qptiff scans.

### Immunohistochemistry (IHC) and Immunofluorescence (IF) quantification

Vectra Polaris IF 40X and Hamamatsu Nanozoomer IHC 40X whole tissue images were uploaded on ImageJ (Fiji) for quantification. Briefly, IF channels were separated into independent images, converted to 8-bit type and tumor area was manually annotated. Next, images belonging to the same channel were equally adjusted to B&W threshold and area fraction, limit to threshold and display labels measurement were set to obtain the percentage of covered area of the signal relative to the tumor area. For IHC, images were first processed with ImageJ plugin IHC toolbox (https://imagej.net/ij/plugins/ihc-toolbox/) using the model H-DAB(browner) to separate the DAB staining from the rest of the tissue. The DAB isolated signal was then processed as the IF to obtain the percentage of covered area of the signal relative to the tumor area.

### RNAscope

RNAscope was performed using the RNAscope Multiplex Fluorescent Reagent Kit v2 (Biotechne, 323110) combined with RNAscope probes against human *HOXD13* (Biotechne, 489291) and *MITF* (Biotechne, 310951) following manufacture instructions. Tissue microarray containing different cutaneous melanoma stages was purchased from tissuearray.com (ME2082d).

### ChIP, ChIPmentation, Hi-ChIP and HiC

Cell lines were double crosslinked with 1% FA (Sigma Aldrich, 47608) and 25uM EGS (Thermo Fisher Scientific, 21565) and processed for ChIP, ChIPmentation and Hi-ChIP.

For ChIP, 1×10^7^ cells were lysed in ChIP lysis buffer (CLB = 10mM Tris HCl pH8, 100mM NaCl, 1mMEDTA pH8, 0.5mM EGTA pH8, 0.1% Sodium Deoxycholate, 0.5% N-lauroylsarcosine, 1% Triton X-100) and sonicated with Diagenode Bioruptor for 15-30 cycles (30” ON +30” OFF max intensity). 5ug/IP of H3K27ac antibody (Active Motif, 91193) or IgG control (Cell Signaling, 3900) was prebound to 50ul of Protein A/G magnetic beads (Pierce Thermo Fisher Scientific, 88803) in PBS+BSA 0.5% for 1hr @RT. Cells’ chromatin was diluted to 1ml in CLB and magnetic beads-antibody complex was added to each IP ON at 4°C on revolver. IP were washed twice in Wash Buffer I (WBI, 20mM Tris HCl pH7.5, 150mM NaCl, 2mM EDTA pH8, 0.1% SDS, 1% Triton X100), Wash Buffer II (WBII, 10mM Tris HCl pH8, 250mM LiCl, 1mM EDTA, 1% Triton X100, 0.7% Sodium Deoxycholate) and TET Buffer (10mM Trist HCl pH8, 1mM EDTA, 0.2% Tween20) and eluted in Elution buffer (EB, 10mM Tris HCl pH8, 300mM NaCl, 5mM EDTA pH8, 0.5% SDS, 10ug RNAse A, 100ug proteinase K) 30min at 37°C + ON at 55°C. IP elutions were purified using MinElute Reaction Cleanup kit (Qiagen, 28204) and libraries were prepared using NEBNext Ultra II DNA Library Prep Kit for Illumina (NEB, E7600S) following manufacture instructions.

For ChIPmentation, the protocol steps are the same as ChIP until the last wash in WBII. Afterward, beads were washed in 10mM Tris-HCl pH8 twice and resuspended in Tagmentation mix (10mM Tris HCl pH8, 5mM MgCl_2_, 10% v/v dimethylformamide) and Tagment DNA Enzyme (Illumina, 20034197) for 10min at 37°C. Next, beads were washed in TET buffer and eluted in EB as described above. ChIPmentation library preparation was done using Schmidl protocol^25^.

For Hi-ChIP and HiC, 1×10^7^ and 2.5×10^6^ cells were processed respectively using Arima-HiC+ kit (Arima Genomics, A101020) following manufacture instructions. Hi-ChIP and HiC libraries were generated using Arima-HiC+ kit (Arima Genomics, A510008, A303011) following manufacture instructions. ChIP, ChIPmentation, Hi-ChIP and HiC libraries were pooled together at 2nM concentration and sequenced paired-end 2×150 using Illumina NovaSeqX+.

### omniATAC

omniATAC protocol was adapted from Howard Chang lab^26^. 5×10^5^ cells were harvested and assessed to not have more than 15% of cell death using trypan blue. Next, dead cells were additionally removed treating the cells with 200U/ml of DNAse (Worthington, LS002007). Cells were resuspended in 500ul of cold buffer 1 (10mM Tris-HCl ph 7.5, 10mM NaCl, 3mM MgCl_2_, 0.1% NP-40, 0.1% Tween-20) and the equivalent volume for 5×10^4^ cells (50ul) was kept for the next steps. A final concentration of 0.1% of digitonin was added to the cells and incubated in ice for 3 minutes. Digitonin permeabilization was quenched adding 1ml cold buffer 2 (10mM Tris-HCl ph 7.5, 10mM NaCl, 3mM MgCl_2_, 0.1% Tween-20). Afterward, cells were pellet and resuspended in 50ul of transposition mixture (20mM Tris HCl pH 7.5, 10mM MgCl_2_, 20% dimethyl formamide, PBS, 1% digitonin, 10% tween-20) supplemented with 2.5-5ul of Transposase (Illumina, 20034197) and incubate in thermomixer at 37°C for 30 minutes with 1000rpm. Transposition mixture was purified using DNA Clean and Concentrator-5 kit (Zymo, D4014) for library preparation. 20ul of transposed DNA was processed for Nextera library using the same protocol described in Schmidl work^25^. omniATAC libraries were pooled together at 2nM concentration and sequenced paired end 2×150 using Illumina NovaSeqX+.

### Cut&Tag

CTCF genomic occupancy was assessed using CUT&Tag-IT^TM^ (Active Motif, 53160) following manufacture instructions. Briefly, 5×10^5^ cells were captured with concanavalin A beads and 1ug of IgG or CTCF antibodies were incubated 2hrs at 4°C on revolver in cell lysis buffer. Afterward, 1ul of guinea pig anti-rabbit antibody was added to the reaction followed by 1hr of tagmentation at room temperature with 1ul of Tn5 transposase enzyme. Tagmentation was stopped with EDTA, SDS and proteinase K incubation for 1hr at 55°C. Tagmented DNA was purified using DNA purification reagents within the kit. Finally, tagmented DNA was processed for Nextera library preparation using i7 and i5 indexed primers and Q5 TAQ polymerase. Cut&Tag libraries were pooled together at 2nM concentration and sequenced paired end 2×50 using Illumina NovaSeqX+.

### ChIP, ChIPmentation, Cut&Tag and ATACseq analysis

For both ChIP and ChIPmentation, demultiplexed fastq files were trimmed and filtered according to the qc using TrimGalore (0.6.6, https://github.com/FelixKrueger/TrimGalore) and fastqc (0.11.7, https://www.bioinformatics.babraham.ac.uk/projects/fastqc/). For ChIP samples, fastq trimming was set for Illumina adaptors; for ChIPmentation, ATACseq and Cut&TAG was set for Nextera adaptors. Alignment to Hg38 human genome was performed using bowtie2 (2.4.1)^27^ and sam files were converted into bam using samtools (1.16)^28^. Bigwig files were generated from bam files using bamcoverage (deeptools 3.5.6)^29^ and filtered out for duplicate and bad genomic regions. Peak calling was performed using MACS2 (2.2.9.1)^30^ using paired-end mode, IgG (ChIP and Cut&TAG) or HA (ChIPmentation) as controls, qvalue 0.05 and both narrowPeak and summit peaks were filtered for blacklisted regions using bedtools (2.31.0)^31^. Peak annotation was performed with homer (4.10)^32^ using narrowPeak files and hg38 genome parameters. DNA motif analysis of CTCF and HOXD13 was done by first expanding of 200bp MACS2 summit peaks with bedtools slop, then fasta sequence was obtained using bedtools getfasta which were processed using MEME-ChIP^33^.

### Hi-ChIP and HiC analyses

For both HiC and Hi-ChIP samples, fastq trimming, qc, alignment, valid pairs identification and matrix generation was performed using HiC-Pro workflow (3.1.0)^34^ using the following parameters: LIGATION_SITE = GATCGATC,GANTGATC,GANTANTC,GATCANTC; BOWTIE2_LOCAL_OPTIONS = --very-sensitive -L 20 --score-min G,20,8 --local –reorder. For HiChIP analysis, H3K27ac loops calling was performed using hichipper (0.7.0)^35^ in which ChIP-seq narrowPeak files of matched samples were used to call and map the loops in the genome. Loops between replicates were merged and filtered using HiCexplorer (3.7.3)^36^. Differential loops analysis was done using diffloop^37^ R (3.5) package. Aggregate Peak Analysis was conducted using GENOVA^38^. For HiC analysis, hic-pro matrixes were converted into juicer and cool formats using hicpro2higlass (HiC-Pro) and hicConvertMatrix (HiCexplorer) respectively. Matrixes normalization was performed using cooler balance (cooltools)^39^ and addNorm (juicer tools)^40^. Compartment analysis was conducted using eigs-cis function (cooltools) and 100kb BIN resolution matrixes. TADs identification was conducted using domaincaller^41^ providing an input matrix of 40kb BIN resolution. Loops were called using both 10kb and 5kb BIN matrixes with Peakachu^42^ and 10kb and 5kb were merged using hicMergeLoops (HiCexplorer). TADs boundaries and loops anchors co-occupied by CTCF were retained for downstream analysis using respectively bedtools intersect and hicValidateLocation (HiCexplorer) respectively. Pyramid plots were generated using GENOVA r package^38^.

### RNA sequencing

Total RNA libraries were prepared using automated TruSeq stranded total RNA and RiboZero Gold kit (Illumina, RS-122-2301) according to the manufacture instructions. RNA libraries were sequenced paired end 2×50 using Illumina NovaSeqX+. Sequencing results were demultiplexed and converted in fastq using Illumina bcl2fastq software. Next, fastq were trimmed and evaluate for qc using TrimGalore (0.6.6) and fastqc (0.11.7). Then, trimmed fastq were aligned to Hg38 human genome and annotated using STAR (2.7.7a)^43^ using –quantMode TranscriptomeSAM GeneCounts and –sjdbGTFfile GENCODEv44. Using R (4.4.3), ReadsPerGene.out.tab were combined in a single table using data.table and processed for normalization and differential expression analysis using respectively edgeR^44^ and DESeq2^45^.

### ABC model

Active-by-Contact (ABC) model^46^ was generated following the authors guidelines (https://github.com/broadinstitute/ABC-Enhancer-Gene-Prediction). Briefly, cCRE were called and measured in activity using makeCandidateRegions and run.neighborhoods scripts in which ATACseq and H3K27ac bam and narrow.Peaks are integrated. Next, HiC matrix were processed using juicebox_dump and compute_powerlaw_fit_from_hic to extrapolate genome wide contacts. ABCscore for each active cCRE was measured using predict function in which cCRE, HiC contacts and RNAseq are integrated. Finally, cCRE were filtered using an ABCscore threshold of 0.02 and self-promoter contacts were excluded for downstream analysis.

### Animal models

For tumor growth assay, 1×10^5^ A375, SKMEL-5, YUMM1.7, B16F10 and D4M-3A cells were injected intradermally in the flanks of 8 weeks old male mice NOD.Cg-Prkdc^scid^ Il2rg^tm1Wjl^/SzJ (NSG) (A375, SKMEL-5 and YUMM1.7) or C57BL/6J (YUMM1.7, D4M-3A, B16F10) (The Jackson Laboratory) in a mix of 50% PBS + 50% Matrigel (Corning, 354234). For the Etrumadenant and Lenvatinib experiment, 8 weeks old male mice C57BL/6J (n=10 per group) were injected intradermally in the flanks with YUMM1.7 HOXD13 and after 3 days of grafting, mice were daily administrated by oral gavage with 10ug/gr of Lenvatinib (MedChem Express, HY-10981) and/or Etrumadenant (MedChem Express, HY-129393) or vehicle (95% corn oil + 5% DMSO).

Every two days, mice were weighed and tumor volume was measured using a two-dimensional caliper and calculated with the formula (*π*/6) × length × wide^47^. All experiments presented in this work respected the maximum tumor volume allowed by NYU Institutional Animal Care and Use Committee (IACUC) of 1500mm^3^. After euthanasia, primary tumors were harvested and fixed in 10% formalin solution for 3 days and stored in 70% ethanol. Fixed tumors were embedded in paraffin, cut and stained by the NYU Langone Health Experimental Pathology core. Animal experiments were approved and conducted under IACUC protocol# IA16-00051.

### Acral Melanoma tissue acquisition, selection, processing, and QC

Formalin-fixed, paraffin-embedded (FFPE) melanoma tissue samples were obtained from the New York University Medical Center, following protocols approved by the Institutional Review Board (IRB H10362). All patients gave their written consent for tissue collection. Blocks were stored at 4° C until use.

FFPE samples were selected for further processing based on artificial intelligence (AI) annotation of tumor area from the digitized clinical hematoxylin and eosin stain (H&E) taken during tissue acquisition. Samples with sufficient tumor area (>5 mm^2^) were selected for preprocessing and quality control.

Each FFPE specimen was cut in 5 µm sections and the first slide was H&E stained. H&E-stained sections were scanned and analyzed for melanoma cells with our in-house generated AI tools^48^. Finally, a 10 µm scroll was cut and used for RNA extraction and RNA quality control. Samples with a DV200 >30% were used for spatial transcriptomics.

### Spatial Transcriptomics Library Preparation and Sequencing

Specimens that passed preprocessing and QC criteria were prepared for spatial transcriptomics following the 10x Genomics Visium/CytAssist protocol for FFPE tissues. Briefly, a tissue section was fixed with methanol, H&E stained and scanned. Next, a 6.5 mm X 6.5 mm region of interest (ROI) was selected and annotated in OMERO Plus (OMERO.web 5.25.0). A clinical pathologist confirmed that melanoma was contained within the designated area. The Visium Spatial Gene Expression Reagent Kit from 10x Genomics was used for permeabilization, reverse transcription, second-strand synthesis, and cDNA amplification. Once finished, pooled DNA libraries were sent for sequencing on Novaseq X+ 10B 100 Cycle Flowcell (Illumina). Reads were mapped against the hg38 human reference transcriptome (GRCh38-2020-A) using Space Ranger v2.1.1 (10x Genomics).

### Analysis of Spatial Sequencing Data

The *filtered_feature_bc_matrix.h5* output files generated by Space Ranger were loaded into Seurat v5.0.1 in an R environment (R v4.3.2 and R Studio v2023.12.0+369). A unique molecular identifiers (UMIs) minimum was used to exclude spots with insufficient RNA or poor capture. Spots with more than 200 UMIs were kept. The Loupe Browser 7.0.1 was used to manually exclude spots that fell outside the tissue borders on H&E. The resultant data were normalized and scaled following the Seurat v5 spatial transcriptomics pipeline. Individual data sets were merged using Seurat after batch correction with Harmony v1.2.

The merged data set was analyzed following the Seurat v5 pipeline. After batch correction with Harmony, a UMAP was established using the first 30 principal components. Data were analyzed at low (0.1) resolution, using canonical melanoma markers (MLANA, S100A1, DCT, SOX10, MITF) to distinguish “Tumor” from “Stroma”. The data sets were then subset to only include clusters identified as “Tumor”. Module scores for activated T-cells (CTL: “CD8A”, “CD8B”, “GZMA”, “GZMB”, “PRF1”, “IFNG”) was generated, added to the Seurat object, and visualized in Violin plots and spatial feature plots.

### Public Datasets mining

#### RNAseq bulk data

To identify the TFs differentially expressed in the Badal dataset, pheatmap was used to perform one minus pearson correlation hierarchical clustering. Graphic representation was generated using ggplot2.

Transcriptomic data and patient clinical information of TCGA cutaneous melanoma samples (SKCM) were downloaded from the cBioportal^49^. Normalized expression counts and integration between TCGA and GTEx and ATACseq profile in TCGA samples were obtained from the UCSC xena browser^50^. H3K27ac ChIPseq fastq files of melanoma STC and NHEM were downloaded from GSE60666 and GSE94488 respectively and processed according to the ENCODE guidelines^51^ to make the tracks comparable to the ENCODE non-melanoma cell lines. HOXD13 expression quantification upon siSOX10 and in different melanoma cell lines were obtained from Scope (https://scope.aertslab.org/#/f98f4e01-5b50-483d-81b5-9ba117343073/*/welcome).

RNAseq fastq files of 91 melanoma patients treated with anti-PD1 or combo anti-PD1 + anti-CTLA4 were downloaded from the SRA BioProject PRJEB23709 and analyzed as described above. MM047 siTEADs RNAseq raw counts were downloaded from GSE60666 and counts per millions (CPM) were calculated using edgeR. Melanoblast, melanocytes and neural-crest HOXD13 expression in human and mouse melanoblast, melanocytes and neural-crest cells was obtained from GSE172069 and GSE140193 respectively.

#### Single cell RNAseq and spatial transcriptomics

Jerby-Arnon^52^, Rambow^53^, Pozniak^54^, Zhang^55^ and Belote^56^ scRNAseq datasets were processed as previously described^57^ and using the authors codes available in their GitHub profiles. Jerby-Arnon, Biermann and Pozniak cell-cell communication prediction and plots were generated using CellChat^58^.

Pozniak and GSE250636 ST Space Ranger outputs are publicly available and were analyzed the same as the acral melanoma samples generated in this study.

### Gene ontology

All the ontological data have been generated using Metascape^59^ and visualized using Prism10.

### Statistical analysis

Statistical test and number of samples are listed in each figure legend. For each experiment at least three biological replicates were used. All statistical analysis were conducted using the most appropriate test in Prism 10 software.

## RESULTS

### The Homeobox-domain transcription factor HOXD13 is upregulated in melanoma compared to naevi and other cancer types

To identify transcription factors (TFs) that could affect melanoma development, we analyzed the expression profiles of a recently updated list of human transcription factors^60^ in a transcriptomic dataset^15^ comprised of cutaneous naevi (n=27) and stage I/II primary melanomas (n=51). Using a one minus Pearson correlation hierarchical clustering approach, we identified 282 TFs significantly upregulated in primary melanoma compared to naevi (Figure S1a, Supplemental Table 1). By ranking their expression across all TCGA tumor samples (n=9,886)^49^ relative to cutaneous melanoma (SKCM) lineage, we identified six TFs strongly associated with SKCM (Figure S1b, Supplemental Table 1). Next, we performed Pearson’s correlation between the expression of the six TFs and signatures of cytotoxic T-cell infiltration (CTL: *CD8A, CD8B, GZMA, GZMB, PRF1, IFNG*) and angiogenesis^61^ (Supplemental Table 1) in TCGA SKCM patients. *HOXD13* was the only TF in this group significantly correlated with increased angiogenesis and reduced CTL (Figure 1a, Supplemental Table 1). HOXD13 was also consistently upregulated in melanoma compared to naevi in other datasets^62^ (Figure S1c). Next, we compared *HOXD13* expression in The Cancer Genome Atlas (TCGA), Genotype Tissue Expression (GTEx; n=17,220)^63^, and the International Cancer Genome Consortium (ICGC; n=8,743)^64^ datasets confirming that *HOXD13* is consistently upregulated in SKCM compared to other cancer types (Figure S1d), uveal melanoma (UVM) and normal tissues (Figure 1b). *HOXD13* expression is also elevated in melanoma compared to other cancer cell lines^65^ (Figure S1e), consistent with its expression in patient tumor tissues. In addition to being overexpressed in primary tumors, we observed a progressive upregulation of *HOXD13* from primary to metastatic melanoma^66^ (Figure 1c, Figure S1f), irrespective of the oncogenic driver^6^ (Figure S1g). RNAscope validates increased *HOXD13* expression in melanoma lesions compared to normal skin on a melanoma tissue microarray (TMA) (Figure 1d, Supplemental Table 2).

**Figure 1:**
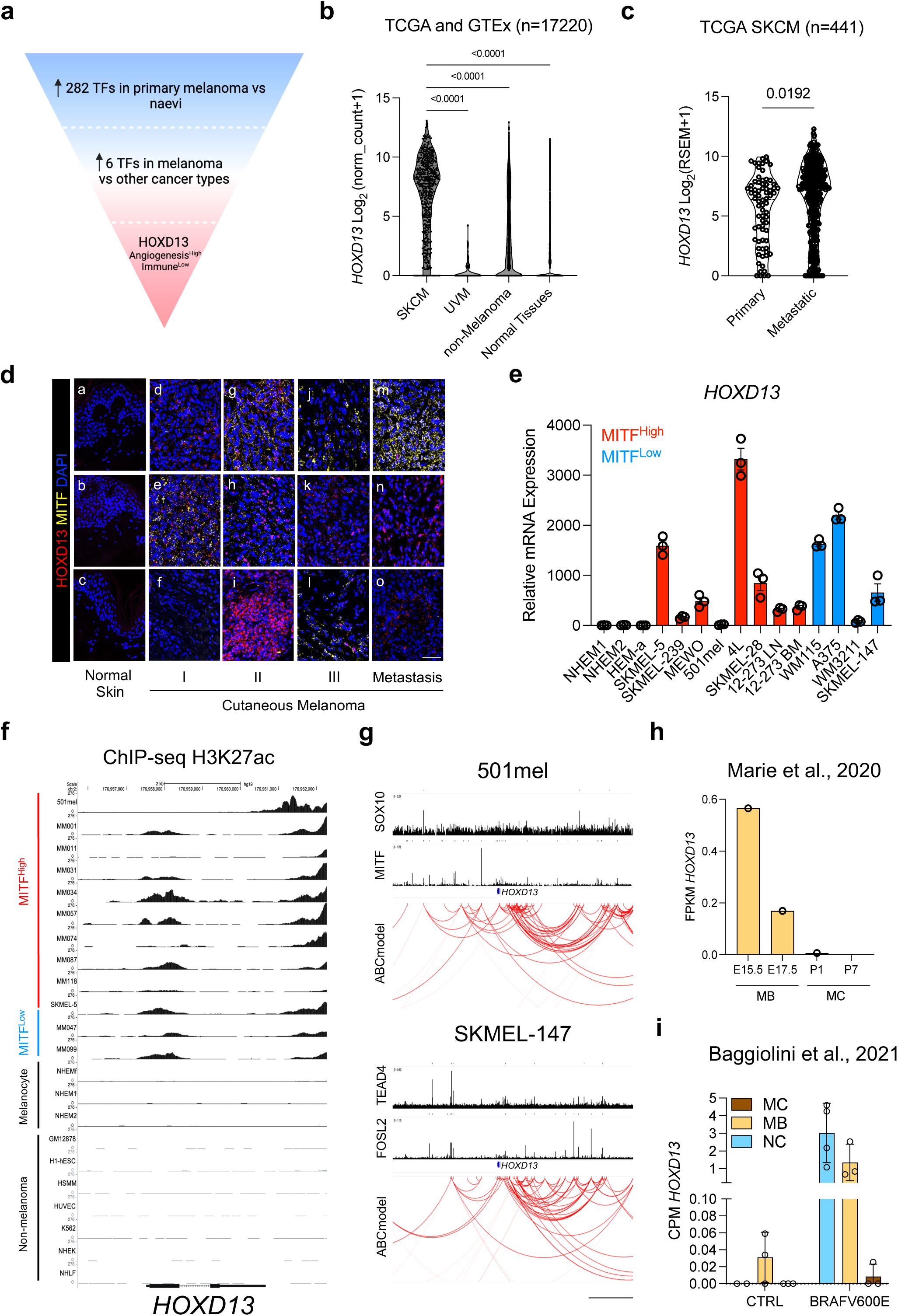
*HOXD13* is upregulated in cutaneous melanoma. **a,** Schematic representation of the analytic approach used to identify HOXD13 as a transcription factor upregulated in melanoma patients with high angiogenesis and low immune infiltration. **b**, Violin plot of *HOXD13* mRNA expression comparing TCGA SKCM patients (n = 469) vs UVM (uveal melanoma, n = 79) vs other cancers (TCGA, n = 9338) and GTEx normal tissues (n = 7414) (p values displayed were obtained using One-way ANOVA Dunnett’s multiple comparisons test). **c**, SKCM patients were segregated into primary (n = 74) and metastatic (367) and *HOXD13* expression is shown in a violin plot. Unpaired t test was performed. **d**, In-situ hybridization for *HOXD13* (red) and *MITF* (yellow) in a TMA containing different skin samples spanning from normal skin (top), primary (stage I, II and III) and metastatic melanoma (bottom) (Supplemental table S1). Black scale bar on the bottom is representative of 200um. **e**, Relative mRNA quantification of *HOXD13* by RT-qPCR in a panel of melanoma cell lines *MITF^High^* (red) and *MITF^Low^* (blue) compared to normal melanocytes (NHEM1, NHEM2, a-HEM). Each dot represents a biological replicate (n = 3 per cell line). **f**, UCSC Genome Browser snapshot of H3K27ac ChIPseq derived from multiple melanoma, melanocytes and non-melanoma cell lines in the *HOXD13* promoter region (marked in sketched black box). H3K27ac deposition is higher in melanoma cells compared to other cell types. **g,** IGV snapshot of the *HOXD13* locus in 501mel and SKMEL-147 melanoma cells showing the genomic occupancy of SOX10, MITF, TEAD4, FOSL2 and the global active regulatory elements contacts generated by the ABC model. Black bar on the bottom represents 200kb size reference. **h.** *HOXD13* expression quantification in murine melanoblasts (MB) and melanocytes (MC) derived from different embryonic stages. **i**, Dot-bar plot showing HOXD13 expression quantification in iPSC-derived neural-crest cells (NC), melanoblasts (MB) and melanocytes (MC) in basal condition or upon ectopic expression of the oncogenic BRAF^V600E^.

We also detected *HOXD13* in melanoma but not stromal cell types in single-cell RNA sequencing (scRNAseq) datasets^52^ (Figure S1h). We found no *HOXD13* enrichment within a specific melanoma cell state including *MITF^High^* (melanocytic, proliferative) or *MITF^Low^*(invasive) melanoma cell states in vivo^53, 54^ (Figure S1i,j) or in vitro^67^ (Figure 1e, Figure S1k). This finding was confirmed on the previously mentioned TMA, where *HOXD13* expression was detected in both *MITF* positive and negative tumors (Figure 1d). Therefore, we hypothesize that HOXD13 is not directly involved in melanoma intra-tumor heterogeneity and that its expression is maintained across melanoma cell states through other gene regulatory networks (GRNs). To address this hypothesis, we analyzed H3K27ac levels in the *HOXD13* locus using ChIPseq data from a large panel of melanoma short-term cultures (STC)^68^, normal human melanocytes^69^ and non-melanoma cells^51^. In line with our previous in silico observations, *HOXD13* promoter was found accessible in both *MITF^High^* and *MITF^Low^* melanoma cells (H3K27ac ChIPseq, Figure 1f) and melanoma patient tissues^70^, with a mean chromatin accessibility directly correlating with *HOXD13* expression (ATACseq vs RNAseq, r=0.409; p<0.0001; Figure S1l). Accordingly, both MITF and SOX10^24^ (characteristic of MITF^High^), and FOSL2/TEADs^69^ (MITF^Low^) TFs bind the regulatory elements contacting the *HOXD13* locus (Figure 1g), and silencing of either SOX10^67^ or TEADs^68^ significantly reduces *HOXD13* expression (Figure S1m). Altogether, these data supports HOXD13 as a TF upregulated in melanoma irrespective of its intra-tumor heterogeneity.

### HOXD13 expression characterizes a melanoblast-like transcriptional program

*HOX* genes are organized in clusters (A-D) and subjected to embryonic positional epigenetic memory, meaning that once the cluster is activated following a transient signal, active chromatin markers are maintained through cell generations^71^. Therefore, we hypothesized that HOXD13 expression in melanoma cells may result from *HOXD* cluster activation during melanocytes’ embryonic development. To test our hypothesis, we mined *HOXD13* expression in melanoblasts (MB) and melanocytes (MC) derived from mouse embryos^72^ and neural-crest cells (NC), MB and MC artificially induced from human induced Pluripotent Stem Cells (iPSCs)^73^ and found HOXD13 being expressed exclusively in MB cells (Figure 1h, Figure S1n). Importantly, introduction of oncogenic BRAF^V600E^ potently induced *HOXD13* expression in NC and MB cells but not in MC (Figure 1i). These data support HOXD13 activation in a malignant context in cells reactivating an MB-like epigenetic program. Accordingly, *HOXD13* expression significantly correlates with a MB gene signature in both SKCM and AM patients (Figure S1o). Altogether, these data support *HOXD13* as a powerful diagnostic melanoma biomarker activated by an oncogenic MB-like epigenetic program, irrespective of intra-tumor heterogeneity and genetic subtypes.

### HOXD13 promotes tumor growth of human and murine melanoma

To gain more insight into the role of HOXD13 in melanoma biology, we inhibited its expression with two independent siRNAs (siHOXD13-1 and siHOXD13-2) validated by RT-qPCR (Figure S2a). *HOXD13* depletion in either *HOXD13*-positive cells (A375) or *HOXD13*-negative 501mel cells had no effect on cell proliferation or cell death in 2D cultures (Figure 2a,b). To assess the role of HOXD13 in melanoma growth in vivo, we stably transduced A375 and SKMEL-5 cells with lentivirus carrying doxycycline-inducible shRNAs^23^ (Fig. S2b) and implanted them subcutaneously into the flanks of NOD/Scid/Il2Rg-/-(NSG) mice. HOXD13 suppression significantly reduced tumor volume (Figure 2c) and weight at termination (Figure S2c). To test whether HOXD13 requirement may be conserved in murine melanoma, we examined *Hoxd13* expression in a panel of murine melanoma cell lines (Figure S2d). Hoxd13 silencing in B16F10 cells recapitulated the growth defect observed in xenograft models (Figure 2d, Figure S2e). These data demonstrate that HOXD13 is necessary for both human and murine melanoma in vivo growth.

**Figure 2:**
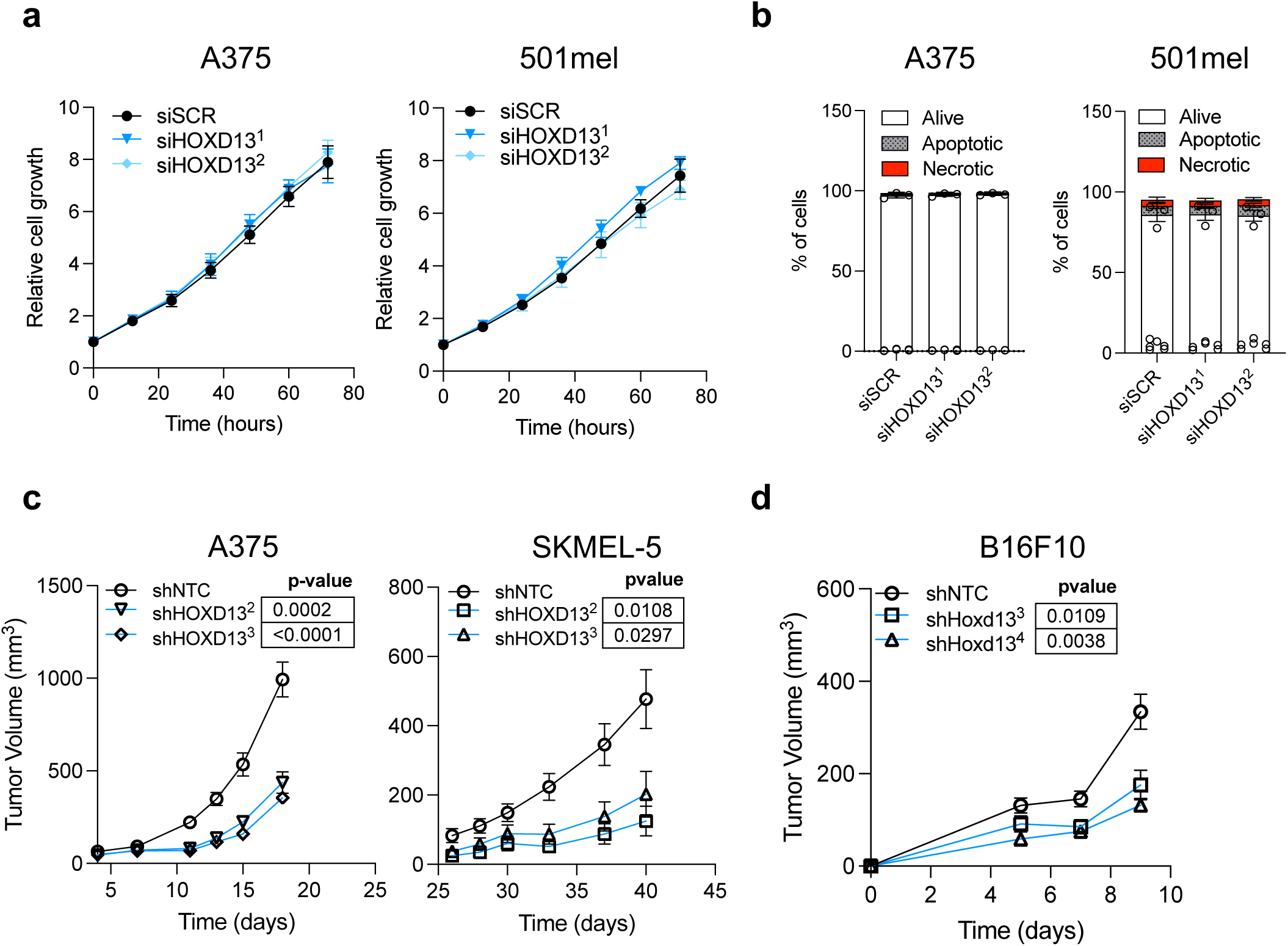
HOXD13 silencing impairs tumor growth. **a**, Growth curves of A375 (left) and 501mel (right) cells upon siRNA control (siSCR) or two independent siHOXD13. Growth was measured every 12hrs for 72hrs. Curves are representative of three biological replicates. **b**, Cell survival of A375 and 501mel cells after 72hrs of *HOXD13* knockdown. No significant differences were observed compared to the control. Alive (AnnexinV-, PI-), apoptotic (AnnexinV+, PI-) and necrotic (AnnexinV+, PI+) cells were detected by FACS. Two-way ANOVA Tukey’s multiple comparisons test was performed. **c**, Tumor growth curve of A375 (left) and SKMEL-5 (right) cell derived xenografts in NSG mouse model. Primary tumors in the flank were measured every 2-3 days. Curves are obtained from 10 biology replicates per condition. ANOVA Tukey’s multiple comparisons test was performed. P values are shown on top right of each curve. **d**, Cell derived syngeneic model of B16F10 murine melanoma cells expressing either a control or two independent shRNAs against *Hoxd1*3. 10 biological replicates were used per shRNA, ANOVA Tukey’s multiple comparisons test was performed. P values are shown on top right of each curve.

### HOXD13 binds distal enhancers and regulates the expression of angiogenesis and immune-modulatory genes

HOXD13 is a master TF of limb development^74–77^. Although HOXD13 has been proposed to be oncogenic in acral melanocytes^78^, its GRN —integrating 3D chromatin contacts and transcriptional output— has not yet been characterized in a cancer context. To do so, we transduced A375 melanoma cells with a N-terminal double-tagged FLAG-HA-HOXD13 doxycycline-inducible construct. ChIPmentation with anti-HA antibodies, identified 29,191 peaks (Figure 3a) enriched with HOX-family DNA binding motif (TTTATDR) (Figure 3b). HOXD13 peak annotation using cis-candidate regulatory elements (cCRE, n= 926,535) defined by the ENCODE project^79^ reveals a preferential binding of this factor to distal enhancers (Figure 3c). To directly link HOXD13 genomic occupancy to its target genes, we performed H3K27ac Hi-ChIP to identify all genome-wide active cCRE contacts (n = 99,374), and mapped HOXD13 binding to loops’ anchors. HOXD13 genomic occupancy greatly overlaps with H3K27ac-mediated loops (n= 16,788/29,191 ∼ 57%) and can be segregated into four clusters based on whether HOXD13 binds the first, second, both, or no cCRE anchors (clusters 1, 2, 3 and 4 respectively; Figure 3d). Intriguingly, cluster 3, characterized by HOXD13 tethering to both anchors, has the most abundant fraction of enhancer-enhancer contacts compared to the other clusters (Figure S3a), showing HOXD13 enrichment at highly interconnected distal regulatory elements.

**Figure 3:**
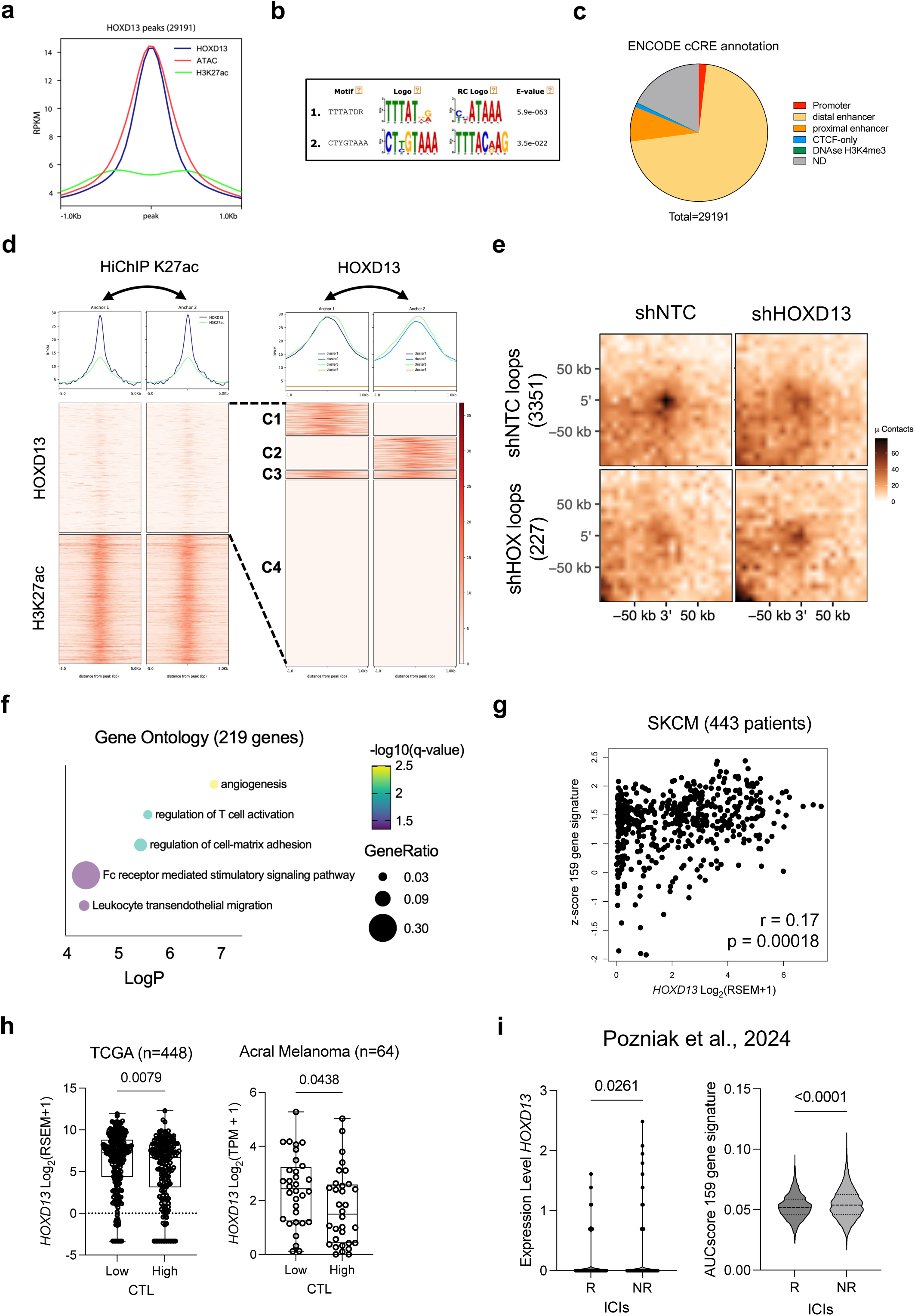
HOXD13 controls immune and angiogenesis genes by establishing 3D contacts between distal regulatory elements. **a**, Genome-wide meta profile of HOXD13, ATACseq and H3K27ac in 29191 HOXD13 peaks identified in A375 cells. A region of +/-1kb around the peak is shown. **b**, MEME-ChIP motif analysis for the 29191 HOXD13 peaks shows an enrichment of HOX binding motif as top hit. **c**, Pie chart shows HOXD13 peak annotation according to the ENCODE project. Most of HOXD13 peaks fall in distal regulatory elements. **d**, (Left) Heatmap profile of the HOXD13 and H3K27ac genomic occupancy on H3K27ac Hi-ChIP anchors (n = 99374). The clustering of these anchors allows to identify 4 anchors clusters differentially bound by HOXD13 (C1, C2, C3 and C4). K-mean clustering was performed using deeptools. **e**, Aggregate Peak Analysis of the significant differential loops (FDR≤0.05) identified with DiffLoop upon silencing of HOXD13. **f**, Multivariable plot representing gene ontology pathways significantly associated to the HOXD13-dependent core of 219 genes. P values, q values and gene ratio were generated with Metascape. **g**, Spearman correlation between 159 genes regulated by HOXD13 and *HOXD13* expression in TCGA melanoma patients (SKCM = 443). Spearman r and p value are shown on bottom right of the plot. **h**, SKCM patients were divided in Cytotoxic T lymphocyte (CTL) low and high based on the expression of 6 genes related to cytotoxic T cell activity. *HOXD1*3 expression is higher in CTL^Low^ patients which are likely to have less T-cell infiltration. Unpaired t test p value is displayed. **i**, Both *HOXD13* and the 159 genes regulated by HOXD13 were scored in a scRNAseq dataset of ICI responder (R) and non-responder (NR) melanoma patients. Unpaired t test p values are shown.

To identify HOXD13-dependent enhancer-promoter looping, we performed H3K27ac Hi-ChIP upon silencing of HOXD13 followed by DiffLoop analysis^37^, which showed a significant loss of cCRE loops (n = 3,351 in control shNTC vs n=277 in shHOXD13) (Figure 3e, Figure S3b,c, Supplemental Table 3). Following RNAseq of shHOXD13 vs control (shNTC-transduced) cells (Figure S3d), we integrated lost and gained loops around gene loci with differentially expressed genes and found a core of 219 genes both deregulated and linked to 3D chromatin loops perturbed by HOXD13 loss (Figure S3e, Supplemental Table 3). Gene ontology analysis of the 219 genes demonstrate that most of them are associated to angiogenesis (i.e., *VEGFA, VEGFB, SEMA3A*) and T-cell modulatory (i.e., *NT5E, CD47, CD70*) pathways (Figure 3f), suggesting that HOXD13 levels may influence the tumor microenvironment^80–85^. Within the HOXD13-dependent 219 genes, we selected those which are downregulated along with *HOXD13* (n=159) as a HOXD13 signature and scored their expression in melanoma patients. This signature correlates with *HOXD13* mRNA levels (Figure 3g) in melanoma tissues, suggesting that HOXD13 also regulates these genes *in vivo*. In line with our ontology analyses, *HOXD13* expression is significantly higher in tumors with low CTL in both cutaneous and acral melanomas (Figure 3h), and also higher in tumors from patients that did not respond to immune checkpoint inhibitors (ICIs) vs responders^52, 54, 86^ (Figure 3i, Figure S3f), both associated with cold or “non-brisk” tumors^87–89^. Altogether, we demonstrate that HOXD13 links highly interconnected enhancers and promotes the expression of genes that might shape the melanoma tumor microenvironment.

### HOXD13 controls a VEGFA/SEMA3A-dependent angiogenic switch

Our functional genomics and transcriptomic data support a role for HOXD13 in regulating genes that modulate angiogenesis which could account for the growth defect observed upon HOXD13 silencing in human xenografts. To identify angiogenesis-related genes consistently affected by HOXD13 perturbation, we performed RNAseq and differential expression analysis in additional murine (HCmel1274, YUMM1.7) and human (SKMEL-5, A375) cell lines subjected to HOXD13 knockdown (KD) by shRNA or overexpression (OE) (Supplemental Table 4). We found *SEMA3A, VEGFA and NT5E* being consistently altered (Figure 4a), with a strong reduction of cCREs’ loops occupied by HOXD13 in their corresponding genomic loci (Figure 4b, Figure S4a). Next, we confirmed that SEMA3A and VEGFA protein levels are deregulated upon silencing or overexpression of HOXD13 by western blot (Figure S4b) or ELISA (Figure 4c). Moreover, the expression of several members of the *SEMA3 and VEGF* pathways significantly correlate with *HOXD13* in melanoma patient tissues (Figure S4c). Next, we mined human melanoma scRNAseq^52, 54, 90^ data to assess whether *HOXD13* expression is associated with angiogenesis pathways directly affecting endothelial/pericyte cells in vivo. Then, we divided malignant cells into high and low *HOXD13* expression (0.1 normalized count cut off) and inferred cell-cell communication pathways in each cell type using CellChat^58^. In line with our previous findings, malignant cells with high *HOXD13* levels display a stronger mediator communication with endothelial/pericyte cells through either VEGF^52, 54^ or SEMA3^52, 54, 90^ pathways (Figure 4d, Figure S4d). In addition, since angiogenesis is an essential regulator of ICIs response^91–93^, we mined both VEGF and SEMA3 pathways activity in *HOXD13^high^* vs *HOXD13^low^*melanoma cells in responder (R) vs non-responder (NR) patients^54^. Here, both VEGF and SEMA3 cell-cell communication between HOXD13^high^ melanoma cells and endothelial cells were predicted to increase in NR patients compared to HOXD13^low^ and R patients (Figure S4e), strengthening the link between HOXD13 activity, angiogenesis, and immune response. To functionally test if HOXD13 modulation impacts angiogenesis in vivo, we examine changes in the vasculature of A375 xenografts and B16F10 syngeneic tumors upon HOXD13 depletion. Indeed, multiplex immunofluorescence (mIF) for CD31 and α-SMA — staining for endothelial and pericyte cells, respectively — revealed a switch in CD31+ and α-SMA+ cells upon HOXD13 KD, suggesting a reduction of α-SMA^High^ and an increase of CD31^High^ vessels (Figure 4e-f, Figure S4f). These data support a crosstalk between HOXD13-expressing melanoma cells and endothelial cells, likely mediated by VEGFA/SEMA3A, that controls an angiogenic switch.

**Figure 4:**
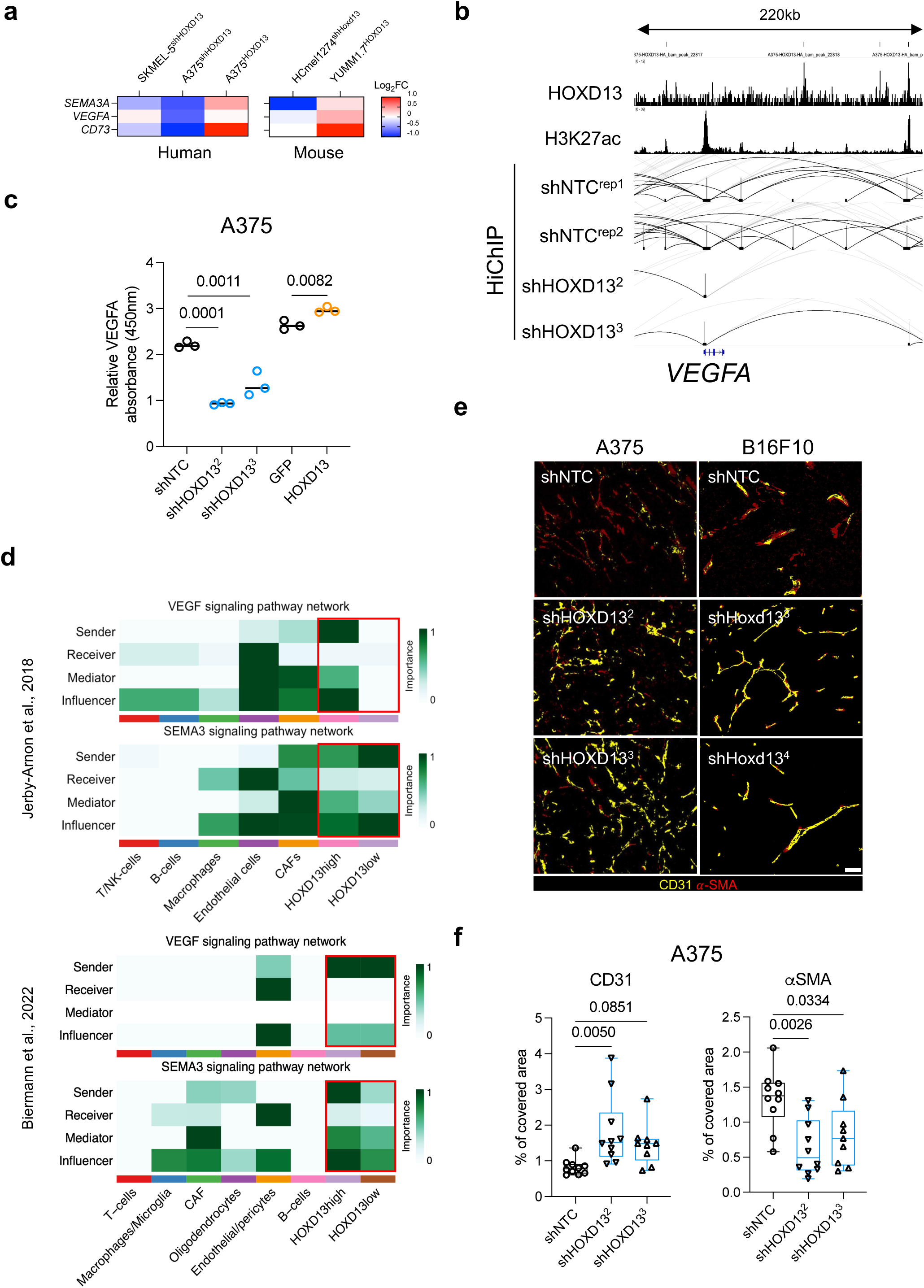
VEGF and SEMA3A pathways are mediated by HOXD13 high expressing cells. **a**, Expression heatmap of SEMA3A and VEGFA in murine (HCmel1274, YUMM1.7) and human (SKMEL-5, A375) melanoma cell lines following silencing or overexpression of HOXD13. Colors’ gradient is based on the Log_2_FC of normalized expression count generated by DESeq2. **b**, Snapshot of *SEMA3A* locus showing the genomic occupancy of HOXD13, H3K27ac and the H3K27ac Hi-ChIP loops in control (shNTC^rep1, rep2^) or following HOXD13 silencing (shHOXD13^2, 3^). Multiple loops are lost upon HOXD13 invalidation which is reflected by the loss of SEMA3A expression. **c**, Secreted VEGF quantification by ELISA shows a significant decrease of VEGFA in cell culture media upon HOXD13 silencing. Conversely, HOXD13 OE led to VEGFA increase. Each dot represents a biological replicate (n=3). One-way ANOVA was performed. Significant p values are displayed. **d**, Cellchat heatmap of the overall importance of cell-cell communication for VEGF and SEMA3 pathway in two independents human scRNAseq datasets. HOXD13 expressing cells are the principal mediators of both pathways. **e**, Multiplex IF for CD31 (endothelial cells) and α-SMA (pericytes) in A375 xenograft and B16 syngeneic tumors control or upon silencing of HOXD13 generates an angiogenic switch unbalancing the ratio of endothelial and pericyte cells. **f**, Box plot quantitation of the overall signal coverage of CD31 and α-SMA in A375 xenograft tumors shows a significant inversion of enrichment of these two markers in shHOXD13 compared to control (shNTC) tumors. One-way ANOVA Dunnett’s multiple comparisons test was performed; p values of the test are shown.

### HOXD13 promotes melanoma immune evasion

The observation of altered immune-related genes upon HOXD13 silencing (Fig. 3f-i) prompted us to test whether HOXD13 regulates T cell function and immune evasion. Therefore, we stably transduced B16F10 and HCMel1274 cells with lentivirus encoding the Ovalbumin antigen (B16F10^OVA^ and HCmel1274^OVA^, respectively) and transfected them with siRNAs against murine Hoxd13. Then, we cultured them in the presence or absence of OVA-activated T cells derived from OT-I transgenic mice. Hoxd13 silencing (Figure S5a) significantly increased T cell-mediated killing in both cell lines (Figure 5a). To test if similar results were observed in a human model, we engineered acute T cell leukemia Jurkat cells to express a T cell receptor (TCR) against NY-ESO peptide, an antigen commonly presented in melanoma cell lines, including A375^94^. HOXD13 silencing (Figure S5b, left) exacerbated A375 growth inhibition in the presence of Jurkat^NY-ESO^ cells (Figure 5b, left); conversely, HOXD13 OE (Figure S5b right) significantly protected melanoma cells in co-culture with Jurkat^NY-ESO^ cells compared to control cells (Figure 5b, right).

**Figure 5:**
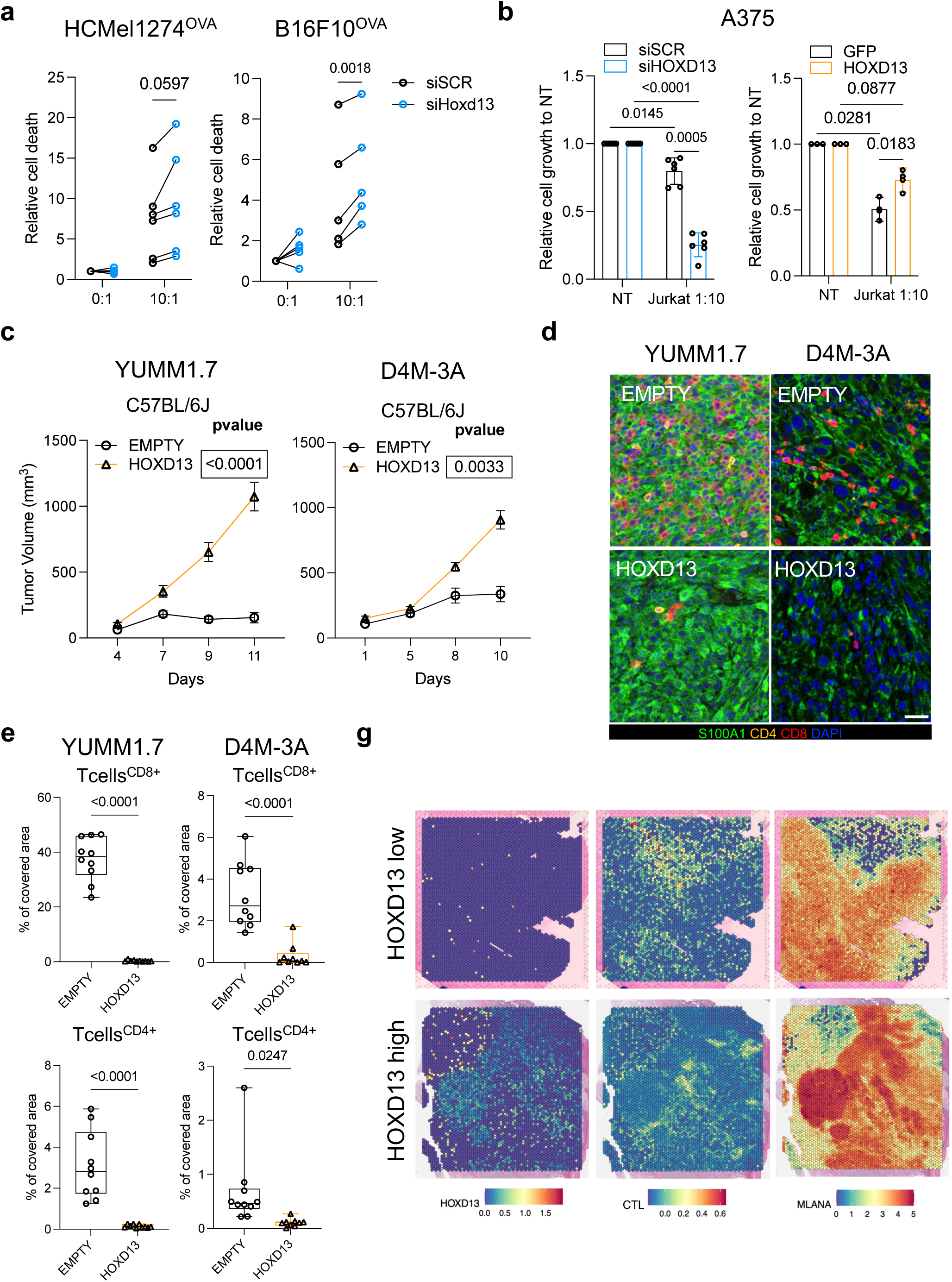
HOXD13 promotes T cell exclusion. **a,** Before-after plot of HCmel1274^OVA^ (left) and B16F10^OVA^ (right) measuring the relative cell death (AnnexinV+ cells) of control and siHoxd13 alone (0:1) or in presence of OT-I T cells (10 times the number of melanoma cells, 10:1). Each dot is a biological replicate, and each line represents a paired experiment. **b**, A375 relative cell growth quantitation in presence or absence of Jurkat^NY-ESO^ and upon HOXD13 silencing (left) or OE (right). Two-Way Tukey’s parametric comparison test was conducted, and p values generated are displayed. **c**, Growth curve of YUMM1.7 and D4M-3A derived tumors in syngeneic animals expressing a control vector (EMPTY) or human HOXD13. HOXD13 ectopic expression boosts tumor growth in syngeneic model. **d**, Representative IF of S100A1, CD4, CD8 and DAPI in YUMM1.7 and D4M-3A derived tumors in syngeneic models. HOXD13 overexpression dramatically decreases CD4 and CD8 T cells infiltration. White bar in the bottom right corresponds to a 50μm scale. **e**, T cells CD8+ (left) and CD4+ (right) quantification in YUMM1.7 and D4M-3A derived tumors based on whole tissue section scanning of 10 biological replicates. Unpaired t test was performed, p values are shown in the dot box plot. **f**, 10x Visium spatial plots of the expression of *HOXD13*, CTL and *MLANA* in two acral melanoma patients bearing low (top) and high (bottom) *HOXD13* expression. Spots counts quantitation was normalized and scaled between the two samples.

Next, we examined if HOXD13 ectopic expression is sufficient to suppress the host anti-tumor immune response and promote the growth of Hoxd13 low cells (Figure S5c). Strikingly, HOXD13 OE dramatically increased tumor growth in immune-competent animals of YUMM1.7 and D4M-3A cells (Figure 5c, Figure S5d,e). mIF for S100A1 (staining cancer cells), CD4 (helper T cells) and CD8 (cytotoxic T cells) revealed a dramatic exclusion of T cells from HOXD13 OE vs control (EMPTY vector) tumors in both YUMM1.7 and D4M-3A models (Figure 5d-e). Finally, spatial transcriptomics of primary acral and cutaneous melanoma samples revealed a significantly lower CTL signature score within the tumor area of HOXD13-expressing acral and cutaneous patients (Figure 5g, Figure S5f). Our data points to HOXD13 as a potent driver of melanoma growth that suppresses both immune infiltration and T cell cytotoxic activity in vitro and in vivo.

### Targeting HOXD13-mediated immune evasion by simultaneously blocking the angiogenesis and adenosine pathways

*NT5E* is a conserved target of HOXD13 across multiple melanoma models (Figure 4a) which encodes for CD73, a membrane-bound 5’-ectonucleotidase that together with CD39, catalyzes the cleavage of extracellular AMP into adenosine. Adenosine is known to bind adenosine A2A receptors (A2AR), suppress immune cells^20, 83, 95, 96^ and modulate vascular permeability^18, 19,97, 98^. We find HOXD13 binds four regulatory elements looping with the *NT5E* promoter and an intronic enhancer, and these 3D contacts are reduced upon *HOXD13* silencing (Figure 6b). In addition, CD73 is significantly upregulated in NR patients and its expression is higher in HOXD13high melanoma cells (Figure S6a). In line with our findings, HOXD13 silencing or OE in both human and murine melanoma cell lines strongly modulate CD73 protein levels, as shown by western blot (Figure 6b) and cytofluorometric analysis (Figure S6b-c). Confirming HOXD13-dependent regulation of CD73 to changes in adenosine extracellular levels, silencing or OE of HOXD13 significantly decreased and increased adenosine levels in culture media, respectively (Figure 6c). Supporting the epistatic relationship of VEGF and CD73 with HOXD13, and a previously reported molecular link between angiogenesis and the adenosine pathway^98, 99^ , combined systemic administration of small molecules targeting angiogenesis (Lenvatinib, VEGFR inhibitor, 10ug/gr) and adenosine signaling (Etrumadenant, A2AR/ A2BR antagonist, 10ug/gr) (combo) significantly reduced the growth-promoting effect of HOXD13 (Figure 6d, Figure S6d). The combo treatment significantly increased immune cell infiltration (Figure 6e), particularly of macrophages, and decreased the tumor vasculature (Figure 6f). These results expose angiogenesis and adenosine pathways as co-operating mediators of HOXD13 pro-oncogenic function in melanoma. In sum, HOXD13 promotes melanoma immune evasion by controlling the tumor vasculature and increasing extracellular adenosine levels, thus simultaneously excluding and inhibiting immune cells’ function. Moreover, our data support the combined targeting of angiogenesis and adenosine pathways as a promising therapeutic approach against HOXD13 expressing melanomas.

**Figure 6:**
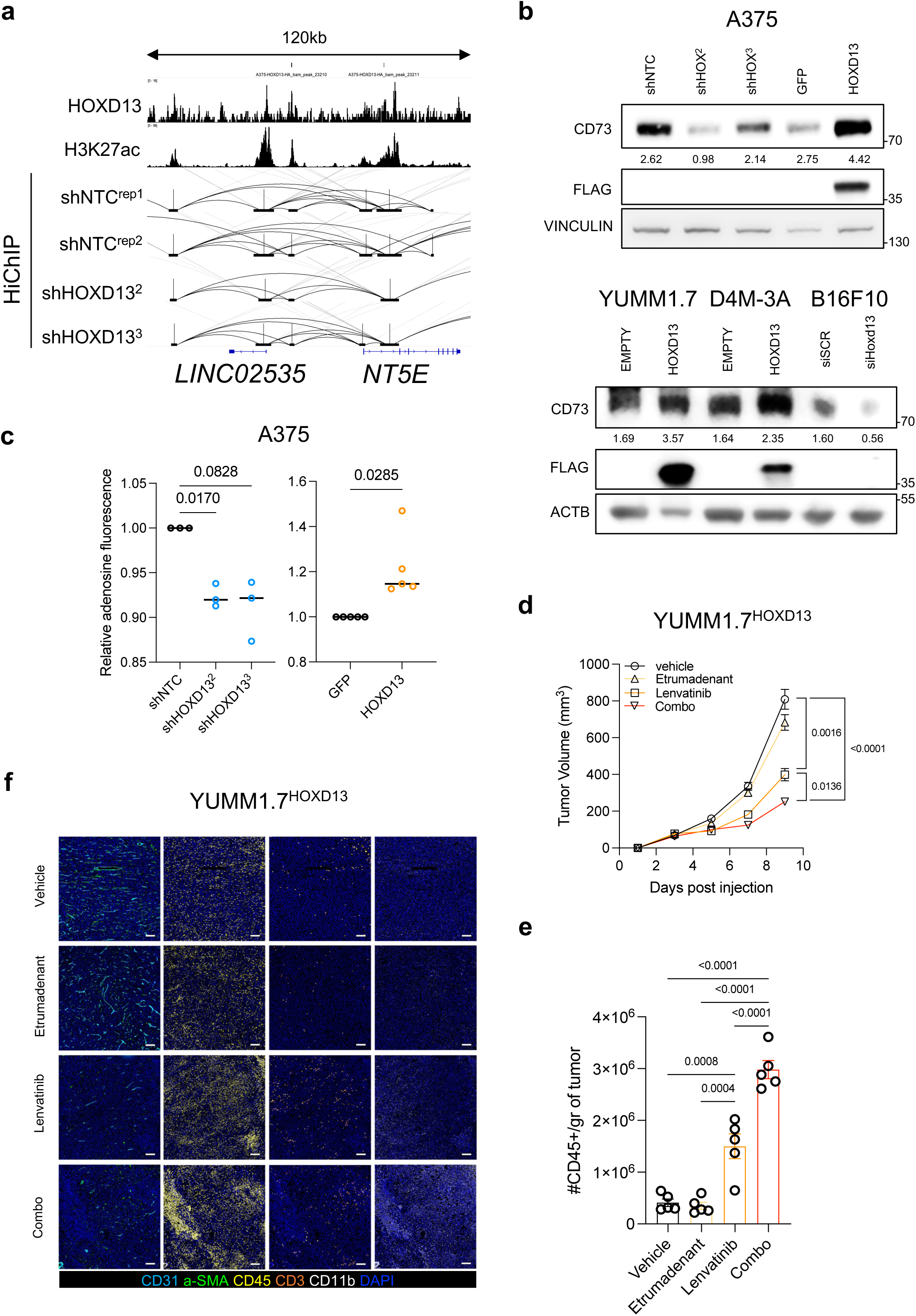
CD73 is a key mediator of HOXD13-dependent T cell exclusion phenotype. **a**, *NT5E* locus displays different regulatory elements contacting each other’s and bound by HOXD13. Silencing of HOXD13 reduces the contacts around *NT5E* promoter. **b**, Representative immunoblot of (*NTF5*-encoded) CD73 modulation by HOXD13 in A375 cells (top) and murine melanoma cells (bottom). Vinculin and actin (ACTB) were used as loading controls. Relative molecular weights are shown on the right of each blot. FLAG stains ectopic expression of HOXD13. **c**, Adenosine was measured in culture media by ELISA in A375 following knockdown (left) or overexpression (right) of HOXD13 which provoked a significant reduction and increase, respectively. One-way ANOVA Dunnett’s multiple comparisons test was used for HOXD13 silencing and paired t test for HOXD13 overexpression. P values are shown in the graphs. **d**, Tumor growth measurements of YUMM1.7^HOXD13^ syngeneic grafts upon vehicle (control), Lenvatinib, Etrumadenant, or Lenvatinib + Etrumadenant (Combo). Combo significantly impairs tumor growth on a synergistic scale compared to the single agents alone. Mixed-effects analysis through Tukey’s multiple comparison was performed and p values are shown in the curves’ graph. **e**. CD45+ cells quantification normalized for the tumors’ weight depicted by flow cytometry. One-way ANOVA non-parametric test was performed; the corresponding significant p-values are shown. **f**, Representative multiplex-IF staining for DAPI, CD45, CD3, CD11b, CD31 and α-SMA in YUMM1.7^HOXD13^ tumors upon vehicle, Etrumadenant, Lenvatinib and combo treatments. Combo dramatically decreased intra-tumor vasculature (CD31, α-SMA) and increased immune cells and, particularly, macrophages infiltration (CD45, CD11b). White bars reflect 500μm size.

## DISCUSSION

Analyzing the expression of hundreds of TFs among all cancer types, we identified HOXD13, whose transcriptional program is expressed in most melanoma metaprograms or states irrespective of their genetic subtype but with high specificity for acral and cutaneous melanoma. We show HOXD13 binds to highly interconnected distal regulatory elements^100^ and those contacts decrease upon HOXD13 silencing, suggesting a role in maintaining multi-enhancer, transcriptionally active hubs, a pioneer behavior reminiscent of many embryonic developmental programs^101^. Among those HOXD13-controlled hubs are its target genes *CD73* and the *SEMA3A/VEGFA* axis whose activity shapes the tumor microenvironment by promoting immune suppression and maintaining an optimal angiogenesis’ balance^95, 102^ (Figure S7).

In previous studies HOX genes’ oncogenic function has been attributed to their effect in proliferation, apoptosis or drug resistance^103, 104^. Despite belonging to an embryonic development program, the molecular triggers of HOX expression in cancer cells are still largely unknown and difficult to link to the tumor cell lineage^105^. Previous studies showed upregulation of the HOX13 subfamily in acral melanoma correlating with melanocyte anatomical location in the limbs’ extremities^78^, the embryonic development program led by HOX13 TFs^77^. However, *HOXD13* expression is not significantly different between volar and non-volar human melanocytes (Figure S8a)^56^. More importantly, *HOXD13* is upregulated in cutaneous malignant melanoma cells compared to acral ones at single cell level (Figure S8b)^55^, suggesting that previous evidence of upregulation in acral melanoma by bulk RNAseq could be explained by expression in non-malignant cells. Like cutaneous melanoma, acral melanomas with high HOXD13 levels display lower immune infiltration, suggesting that HOXD13 oncogenic function may be conserved between cutaneous and acral lesions. During melanocytes’ development, HOXD13 is expressed exclusively in MB cells and melanoma cells may reactivate *HOXD13* through a complex regulation by SOX10 and TEADs. In addition, during limb development, HOXD13 expression in the cells of the primordial limbs is regulated by a dynamic shift between two Topological Associated Domains (TADs) boundaries, C-DOM and T-DOM, which repress and activate *HOXD13* to commit the cells toward the formation of the limb or the forearm^76^, respectively. Integrating HiC, ATACseq, CTCF CUT&Tag and RNAseq, we compared the TADs organization around the *HOXD13* locus in 501mel (*HOXD13* negative) vs SKMEL-5 (*HOXD13* positive) and found that *HOXD13* falls in the boundary between the two TADs resembling the C-DOM and T-DOM (Figure S8c). Specifically, the *HOXD13* locus lies on the T-DOM-like TAD in 501mel and in the C-DOM-like TAD in SKMEL-5 which regulate the recruitment of regulatory elements from the C-DOM-like TAD contacting and activating HOXD13 as demonstrated by the ABC model (Figure S8d). However, since melanoma UV-dependent C>T/CC>TT substitution can impair CTCF genomic occupancy^106, 107^, we cannot exclude an aberrant CTCF repositioning at C-DOM and T-DOM boundaries leading to HOXD13 hyperactivation.

Immune evasion is one of the hallmarks of cancer^108^, particularly in the early phases of melanoma formation in which high rate of UV somatic mutations promote neoantigen formation and immunoediting^109^. Therefore, melanoma cells must immediately find ways to avoid the host immune response. Different immunosuppressive genetic mechanisms have been described which range from point mutations to broad chromosomal rearrangements^110, 111^ involving key immune-related genes such as PD-L1/2^112^, major complex of histocompatibility class I and II (MHCI and MHCII)^113, 114^ and IFNγ^115^, but also oncogenic pathways such as WNT^116, 117^, PI3K^118^, CDK4/6^52^ and Mitogen-Activated Protein Kinase (MAPK)^13^. In parallel, there has been growing evidence of non-mutational immunosuppressive mechanisms involving GRNs dysregulation^54, 119, 120^ and epigenetic reprogramming^121, 122^. Here, we identified a novel epigenetic, HOXD13-dependent immunosuppressive pathway activating CD73 that increases extracellular adenosine and impairs T cells immune surveillance. Previous work has shown that CD73 is upregulated by TGFβ in dedifferentiated melanoma cells upon nutrient depletion, resulting in increased extracellular adenosine levels^123^ which also correlate with MAPK inhibitors resistance^124^. Curiously, we didn’t observe EMT-like or TGFβ-related genes deregulated upon HOXD13 perturbation which prompts us to hypothesize that, in the absence of HOXD13, CD73 as well as VEGF and SEMA3A are regulated by other pathways and GRNs (e.g., TGFβ). Moreover, CD73 has been shown to be secreted^124^, providing another level of complexity in CD73 regulation and a larger impact beyond the tumor-mediated effect in boosting adenosine levels in the stromal microenvironment.

Angiogenesis —the formation of new branches of blood vessels from pre-existing ones— is considered an essential hallmark of cancer^108^ since it ensures a continuous supply of nutrients and oxygen levels to a dysplastic mass in perpetual growth; meanwhile, new vessels can be conduits for anti-tumor immune surveillance. Unbalanced endothelial and pericytic cell growth can lead to leaky and permeable vessels which may less functional^125–128^. Recent views of angiogenesis suggest that an optimal balance of pro- and anti-angiogenic stimuli from both cancer and stromal cells^129^ including the two antagonistic pathways VEGF (pro-angiogenic)^102^ and SEMA3A (anti-angiogenic)^130^ orchestrate an “angiogenic-switch” and establish an endothelial to pericyte cell ratio. Consistently, while there is no evident correlation between vascular density and melanoma patient outcomes^11, 131^, levels of angiogenic-switch factors and particularly the VEGF/SEMA3A ratio^132^ hold strong prognostic value^133–135^. Our in vitro and in vivo functional assays and in silico analysis in three independent scRNAseq datasets^52, 90^ suggest that modulation of pericyte: endothelial cell ratio by HOXD13 not only promotes tumor normoxia^136^ and cancer cell progression^137, 138^, but may also potentially decrease response to ICIs which will be addressed in future studies. The therapeutic value of targeting simultaneously angiogenesis and immune response is currently being explored on ongoing clinical trials combining ICIs with angiogenesis inhibitors^92, 93^, or using bispecific anti-PD1/VEGF antibodies^139^.

Since direct HOXD13 targeting is difficult, patients’ stratification based on HOXD13 expression could help identify ICI-resistant patients who could benefit from a combinatorial treatment against HOXD13 targets, adenosine (ADO), and angiogenesis pathways. Inhibitors of the ADO pathway, such as promising anti-CD73 antibodies (e.g. NCT05431270, NCT04672434, NCT04322006, NCT04572152) and A2R small molecule inhibitors (e.g. NCT04262856, NCT04660812, NCT04969315, NCT03846310) are currently being tested in clinical trials, alone or in combination with ICI^140^. Our data suggests that patients with HOXD13+ tumors can benefit from these treatments, alone or in combination with anti-angiogenic factors, such as SEMA3A-^141^ or VEGF-blocking antibodies, or Lenvatinib, a pan-tyrosine kinase inhibitor (TKI) particularly effective against VEGF receptors^142^. Given the observed conservation of the HOXD13-angiogenesis-adenosine axis between cutaneous and acral melanoma, our findings open a new therapeutic opportunity for acral tumors known as ICI refractory.

## Supporting information

Supplemental reagents

## AUTHOR CONTRIBUTION

P.B. and E.H conceived and designed the study. P.B., A.F.Y. performed and F.R. helped with in vivo experiments. C.D. and J.S. provided guidance with the Hi-ChIP analysis. I.W.S. generated the TFs ranking in TCGA. E.V. and C.D.R.E provide transcriptomic analysis of acral melanoma from a Mexican cohort. T.S. helped with the ABC model analysis. I.D. performed immune cells’ flow cytometry and analysis with the supervision of A.L. R.S. and M.S. performed spatial transcriptomic analysis on patients’ samples provided by I.O. P.B. performed most of the experiments in vitro and in vivo. T.S. helped in setting the ABCmodel. P.B. performed scRNAseq, ST, ChIPmentation, ATAC, Cut&Tag, HiC, Hi-ChIP, RNAseq analysis. P.B. and E.H. interpreted the data and wrote the manuscript.

## AKNOWLEDGMENTS

We thank members of the Experimental Pathology Research Laboratory (Director: Dr. Cindy Loomis) for tissue processing and histology staining and the Genome Technology Center (Director: Dr. Adriana Heguy) at NYULH for RNA library preparation and sequencing. These core facilities are partially funded by the Cancer Center Support Grant (NCI/NIH; 5P30CA016087) to NYU Perlmutter Cancer Center. We thank Dr. Giulia Cova (Skok lab) for sharing the omniATAC and ChIPmentation protocols. This work was funded by the NIH/NCI R01CA274100 (MPI: E.H., M.S.), and U54 CA263001 (MPI: E.H., I.O.). P.B. has been funded by the National Cancer Center, Melanoma Research Alliance and Melanoma Research Foundation fellowships.

## Supplemental Figures

**Figure S1:**
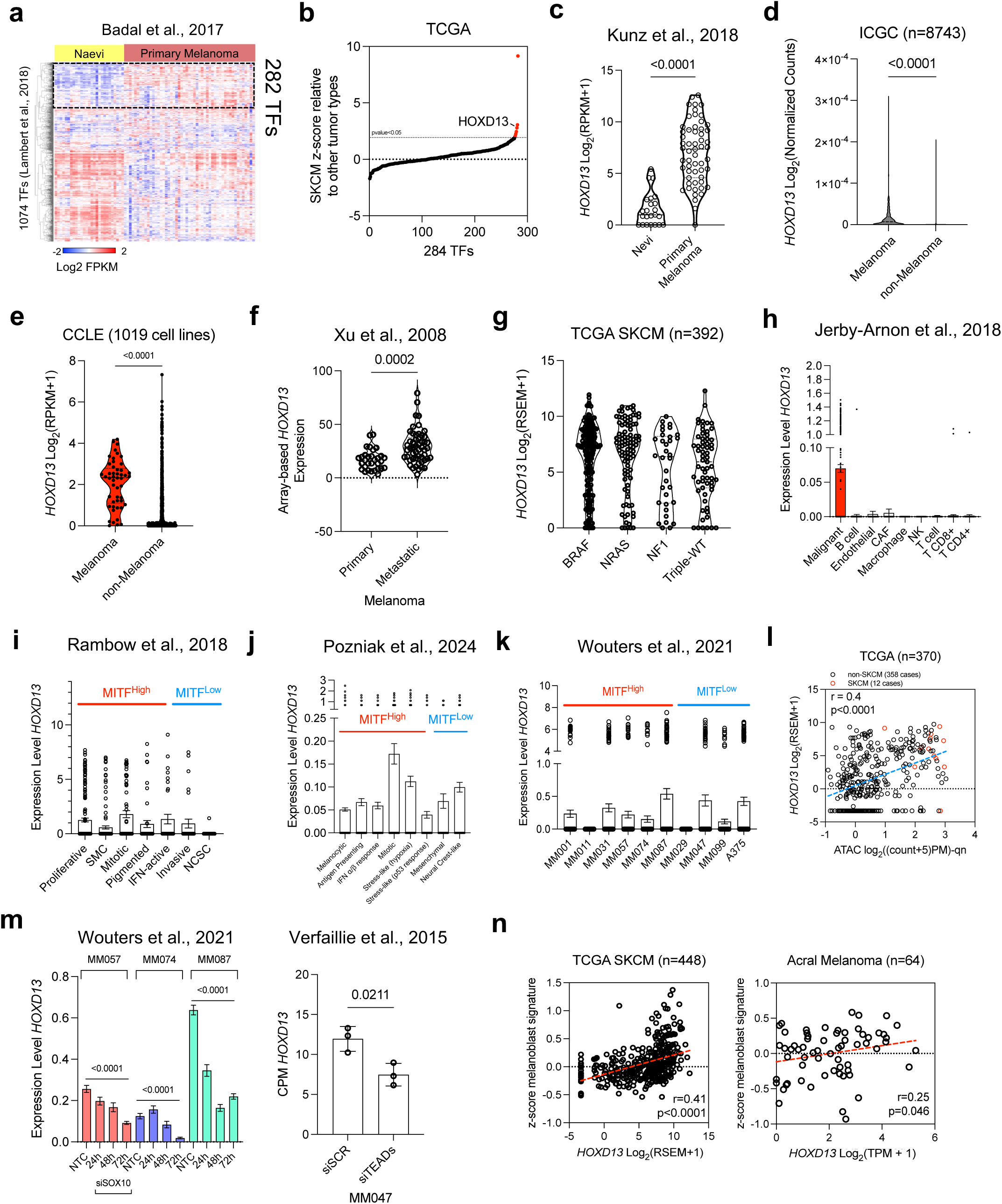
**a**, Heatmap representing the hierarchical clustering of Log_2_FPKM expression of 1074 TFs annotated according to Lambert et al., 2018 between naevi (n = 21) and primary melanomas (n = 57). 282 TFs significantly upregulated in primary melanomas are highlighted on the top right. **b**, A dot plot showing the z-score ranking of 282 TFs (previously identified as upregulated in primary melanoma vs naevi) in all TCGA patient samples (n = 9886). In red are marked the 6 TFs whose p value is less than 0.05 according to the z-score normal distribution relative to SKCM patients. **c**, *HOXD13* mRNA expression in Kunz et al., 2018 is significantly higher in Primary melanoma vs naevi. P value was generated by unpaired t test. **d**, Violin plot representing the significant upregulation of *HOXD13* in melanoma vs other types of cancer in the ICGC consortium (n = 8743). Unpaired t test was performed, p value is displayed. **e**, *HOXD13* expression is higher in melanoma cell lines compared to other cancer cell lines (CCLE, n = 1019). Unpaired t test was conducted to generate the displayed p value. **f**, Violin plot showing the microarray-based expression of *HOXD13* in primary and metastatic melanoma samples. As observed in TCGA, *HOXD13* is upregulated in metastasis vs primary melanoma. P value was generated with an unpaired t test. **g**, *HOXD13* expression is not significantly associated to any melanoma genetic subtype (BRAF, NRAS, NF1 or Triple-WT). **h**, Dot-bar plot representing *HOXD13* expression at single cell level (n = 7186) in different cell types in human melanoma patients (n = 31) generated by Jerby-Arnon et al., 2018. *HOXD13* is mostly expressed in the malignant compartment. **I**-**j**, Dot-bar plot showing *HOXD13* transcript levels across melanoma cell states annotated in two independent scRNAseq datasets. MITF^High^ and MITF^Low^ are marked based on the authors conclusions. **k**, scRNAseq expression levels of *HOXD13* in multiple melanoma cell lines belonging to different states marked by different colors. **l**, Dot plot correlation between *HOXD13* expression and the average counts of ATAC signal in the *HOXD13* promoter derived from multiple cancer patients. In red are marked cutaneous melanoma patients which are skewed on the top right, suggesting a higher promoter activity and expression of *HOXD13* compared to other non-melanoma patients. Spearman correlation r is shown on the top left. In sketched blue is shown the simple linear regression line. P value<0.0001. **m**, (left) scRNAseq *HOXD13* expression in three melanoma cell lines is significantly downregulated upon silencing of SOX10 (MM057 n = 8963, MM074 n = 7255, MM087 n = 8131). One-way ANOVA Dunnett’s multiple comparisons test was done within each cell line. (Right) Box plot representing *HOXD13* mRNA significant downregulation following the silencing of all TEADs TF. Black dots represent biological replicates. Unpaired t test was performed. P value is shown. **n**, Dot plot representing the positive, significant correlation between *HOXD13* and a melanoblast signature (n=45 genes) in both cutaneous (top) and acral (bottom) melanoma patients.

**Figure S2:**
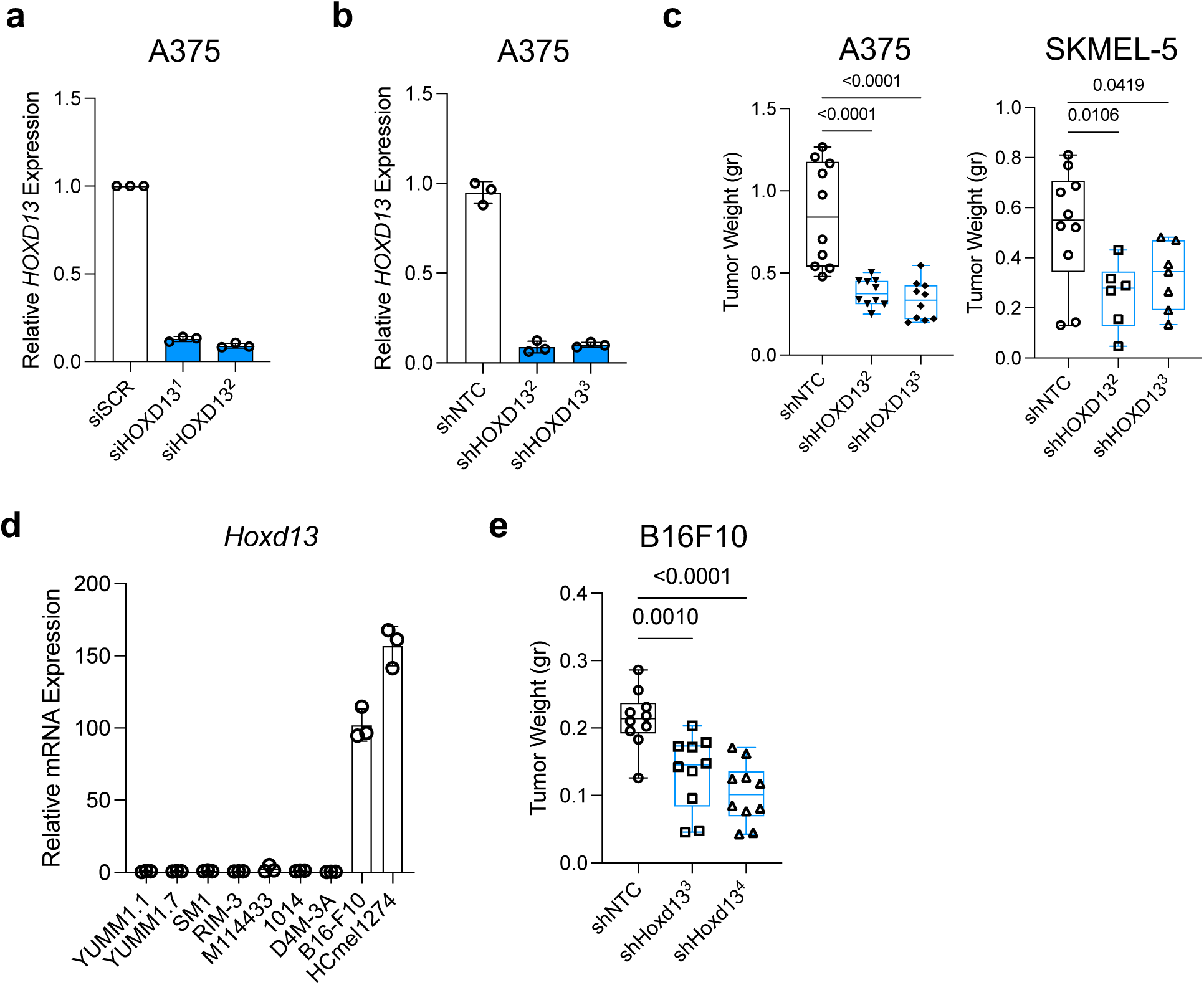
**a**, Relative mRNA quantification of *HOXD13* in A375 in siSCR (control) and upon two independent siRNAs against HOXD13. Each dot represents a biological replicate (n = 3). **b**, Bar-dot plot representing *HOXD13* mRNA downregulation using two different doxycycline inducible shRNAs compared to a control (shNTC). Biological triplicates are represented by each dot (n = 3). **c**, Box-dot plots of A375 (left) and SKMEL-5 (right) derived xenograft tumors’ weight harvested after induction of shRNA control (shNTC) or against *HOXD1*3. P values displayed were generated using a one-way ANOVA Dunnett’s multiple comparisons test. **d**, Murine *Hoxd13* expression was measured in a panel of mouse melanoma cell lines. B16F10 and HCmel1274 were the cell lines showing greater expression of Hoxd13. **e**, B16F10 syngeneic tumors have a reduced weight upon Hoxd13 invalidation. One-way ANOVA Dunnett’s multiple comparisons test was performed; p values are displayed above the box-dot plots.

**Figure S3:**
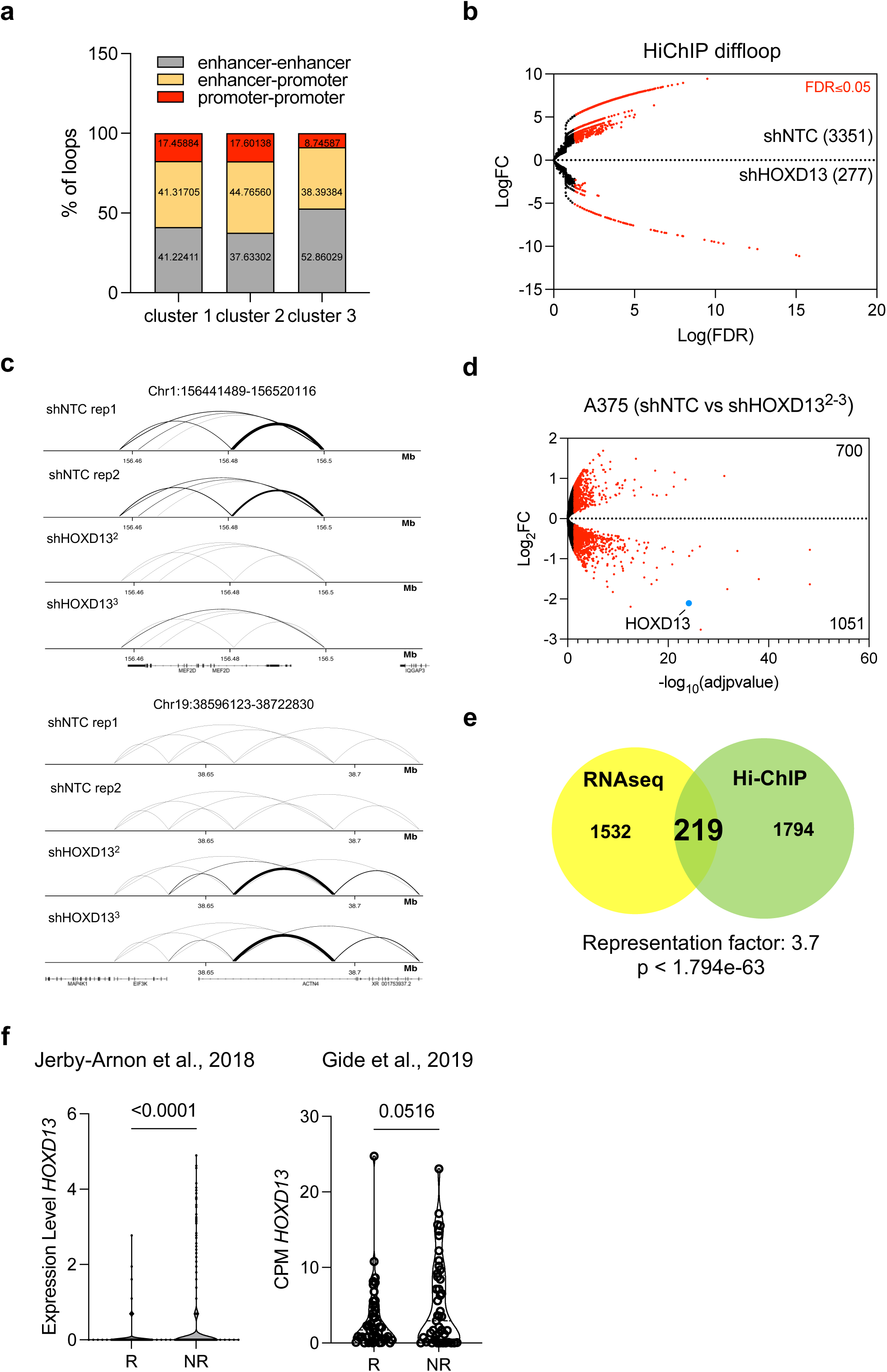
**a**, Grouped histogram showing the percentage of enhancer-enhancer, enhancer-promoter and promoter-promoter loop interactions across clusters 1, 2 and 3 identified by k-mean clustering of H3K27ac Hi-ChIP co-occupied by HOXD13. **b**, Volcano plot of all H3K27ac Hi-ChIP differential loops between shNTC and shHOXD13 A375 cells based on the LogFC and the LogFDR. In red are marked the loops with an FDR≤0.05. Multiple peaks are lost in shHOXD13 (n = 3351) and few are gained (n = 227). **c**, Diffloop snapshot of two additional examples of loops gained and lost upon silencing of *HOXD13*. **d**, Volcano plot representing all the differentially expressed genes in A375 between shNTC and shHOXD13. In red are highlighted the genes with an adjusted pvalue≤0.05. In blue is marked HOXD13. In total, 700 genes are upregulated and 1051 are downregulated when HOXD13 is knockdown. **e**, Venn Diagram showing the integration between differentially expressed genes identified by RNAseq and the genes associated to the differential loops identified by Diffloop in A375 cells. A core of 219 genes significantly overlaps. Hypergeometric test representation factor and p value are displayed. **f**, *HOXD13* expression measured in (left) Jerby-Arnon scRNAseq dataset and (right) Gide bulk RNAseq cohort (n = 91) comparing responder (R) vs non-responder (NR) to immune checkpoint inhibitors (ICIs) shows a significant upregulation of *HOXD13* in NR patients. P values are shown and generated following unpaired t test.

**Figure S4:**
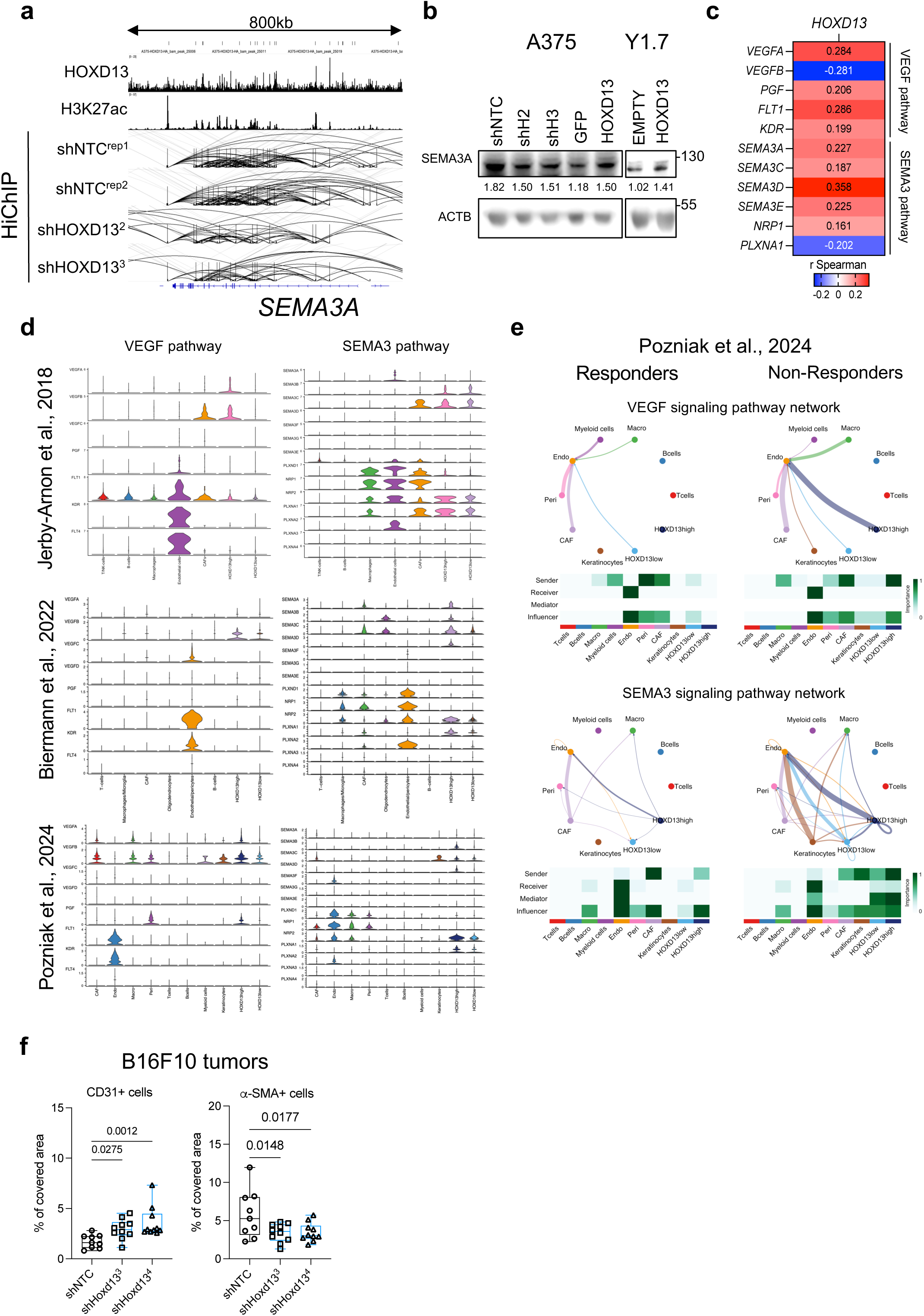
**a**, Similar to *SEMA3A, VEGFA* locus is bound by HOXD13 in different regulatory elements and HOXD13 loss reduces the number of H3K27ac loops around this region. **b**, Immunoblot of SEMA3A in human (A375) and murine (YUMM1.7 = Y1.7) melanoma cell lines upon knockdown or overexpression of HOXD13. β-actin (ACTB) is shown as loading control. **c**, Spearman correlation heatmap between HOXD13 expression and major members of the VEGF/SEMA3 pathways derived from SKCM TCGA patients (n = 443). **d**, Violin plots generated using CellChat showing the expression of the main ligand, receptors and effectors of VEFG and SEMA3 pathways in different cell types annotated in Jerby-Arnon, Biermann and Pozniak datasets. Malignant cells *HOXD13* high have a greater enrichment of both VEGF and SEMA3 components compared to *HOXD13* low. **e**, CellChat circle plot and heatmaps of VEGF and SEMA3 cell-cell communication strength signaling in R and NR patients. VEGF and SEMA3 signaling dramatically increase in NR patients between *HOXD13^high^* malignant cells and endothelial cells. **f**, Box-plot quantification of the percentage of covered area of CD31 and α-SMA IF signal in B16F10 syngeneic tumors control (shNTC) and knockdown for Hoxd13 (shHoxd13^3^, shHoxd13^4^). One-way ANOVA test was performed. Significant p values are shown in the graphs.

**Figure S5:**
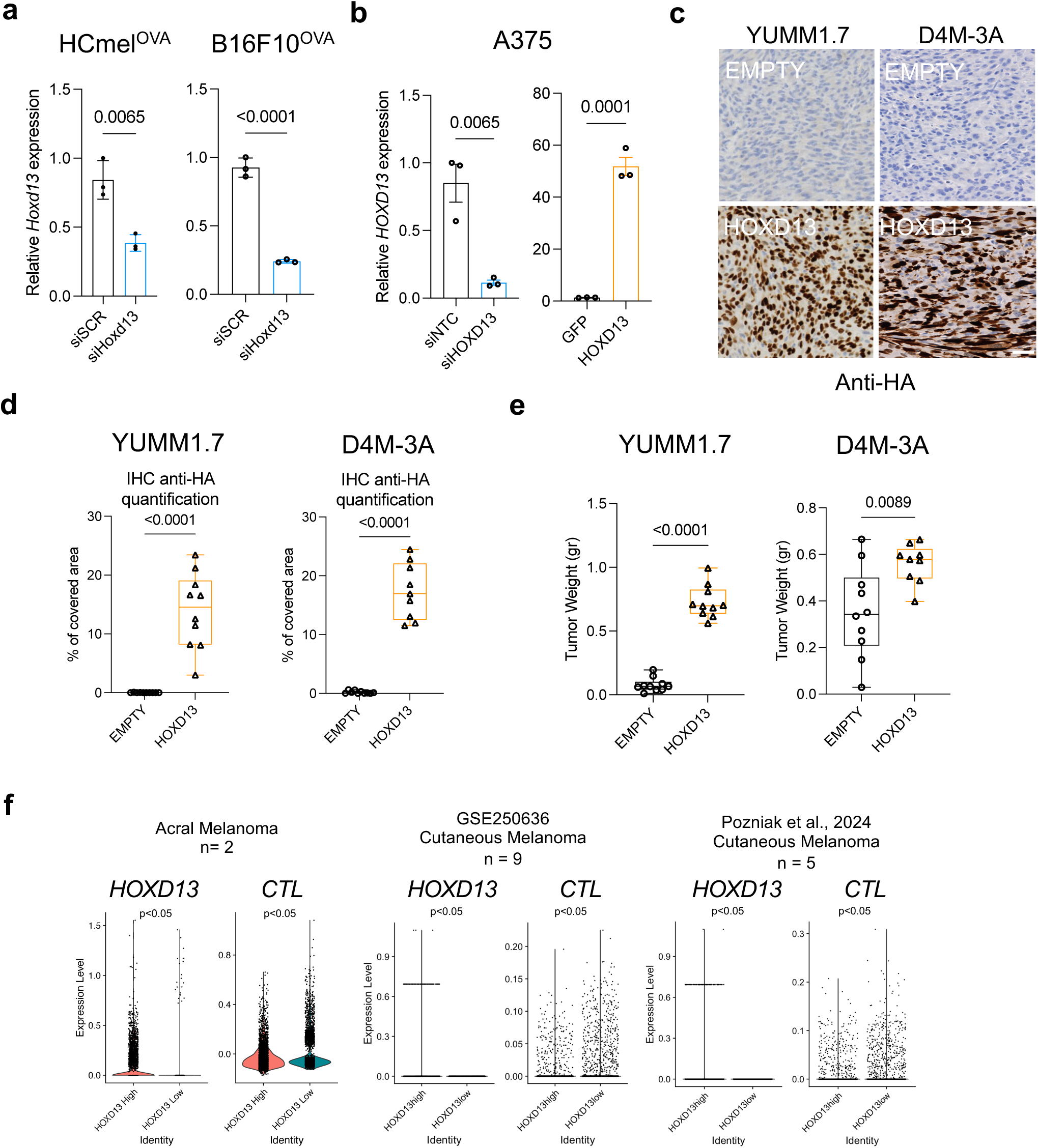
**a**, RT-qPCR for Hoxd13 in HCmel1274^OVA^ and B16F10^OVA^ following siRNA treatment. Unpaired t-test p values are shown using three biological replicates. **b**, mRNA quantification by RT-qPCR of HOXD13 in A375^RFP^ cells upon knockdown (left) or OE (right). Unpaired t-test was conducted, p values are shown. **c**, IHC representative of the anti-HA staining in YUMM1.7 and D4M-3A derived tumors in C57BL/6J animals. HA staining marks HOXD13 protein which is strongly enriched in HOXD13 overexpressing tumors compared to the EMPTY vector control. White bar in the bottom right correspond to a 50μm scale. **d**, Dot-box plot showing the HA staining quantitation in YUMM1.7 and D4M-3A derived tumors in immunocompetent animals. 10 independent biological replicates per group were collected (EMPTY vs HOXD13). Unpaired t test was conducted, p values are shown on top of the dot-boxes. **e**, Weight measurement of YUMM1.7 (left) and D4M-3A (right) tumors derived from syngeneic models. Each dot and triangle represent a biological replicate (n = 10 per group). P values were obtained performing unpaired t test. **f**, Spatial transcriptomics violin plots showing the quantification of HOXD13 (left) and CTL signature score (right) in malignant spatial areas of acral (n=2) and cutaneous (GSE250636 n=9; Pozniak n=5) melanoma paraffin tissue sections. Unpaired t-test was performed, and p values are shown in the graphs.

**Figure S6:**
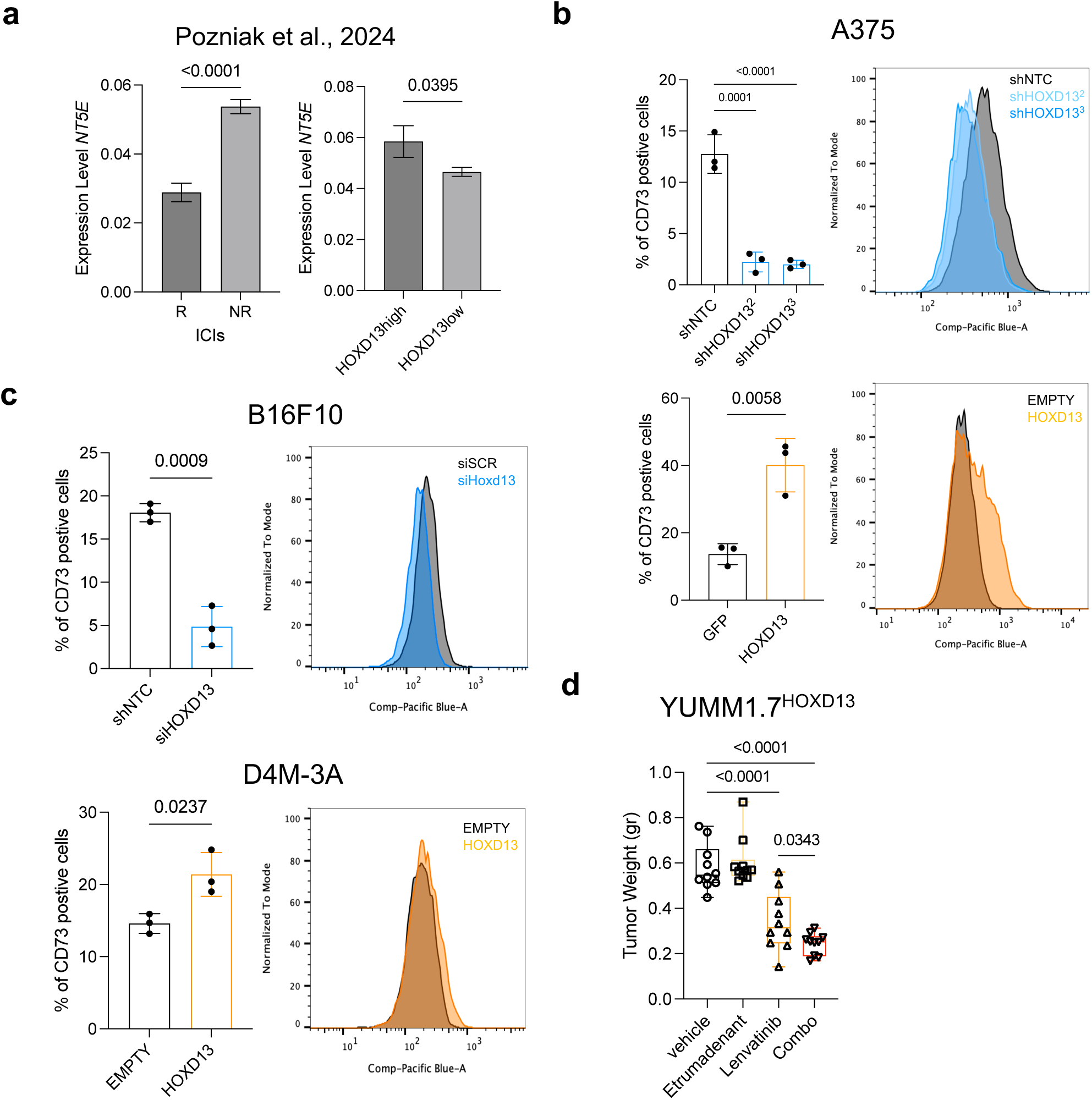
**a**, CD73 (*NT5E*) scRNAseq expression quantitation in R and NR (left), and in HOXD13^high^ vs low malignant cells. Unpaired t-test p values are shown. **b-c**, CD73 surface protein was measured by FACS in A375, D4M-3A and B16F10 cells overexpressing or silencing HOXD13. (Left) Dot-bar plot showing the percentage of CD73 positive cells based on an arbitrary threshold. (Right) A representative profile of CD73 stained cells shifting upon HOXD13 perturbation. Each dot represents a biological replicate (n = 3). One-way ANOVA Dunnett’s multiple comparisons test was performed; P values are shown. **d**, Box-plot YUMM1.7^HOXD13^ tumor weight quantification from mice treated with vehicle (control, n=10)), Lenvatinib (n=10), Etrumadenant (n=10) or Combo (Lenvatinib + Etrumadenant, n=10). One-way ANOVA Dunnett’s multiple comparisons test p values are displayed.

**Figure S7:**
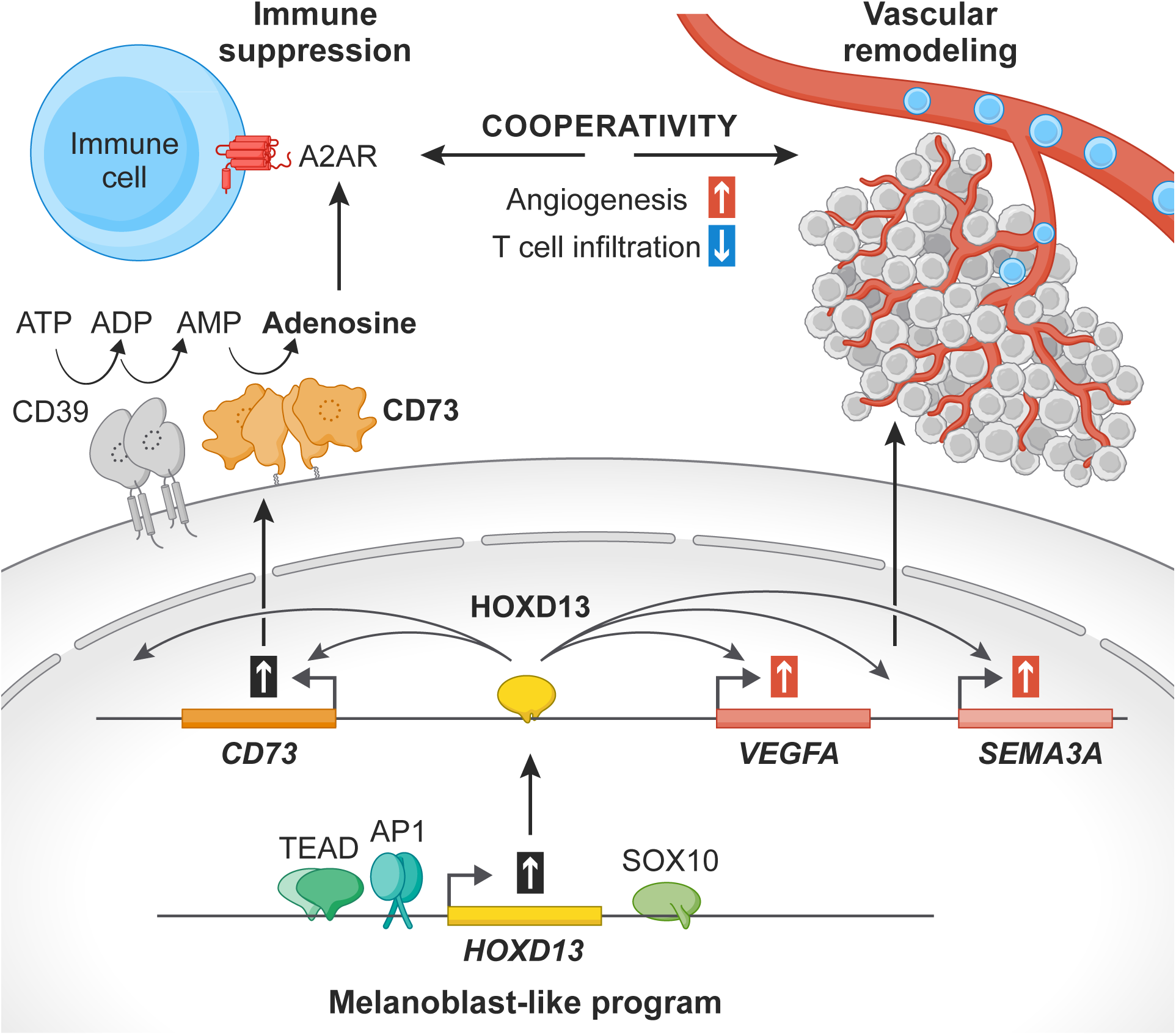
Working model of the HOXD13 gene regulatory network in melanoma cell simultaneously driving immune suppression and vascular remodeling.

**Figure S8:**
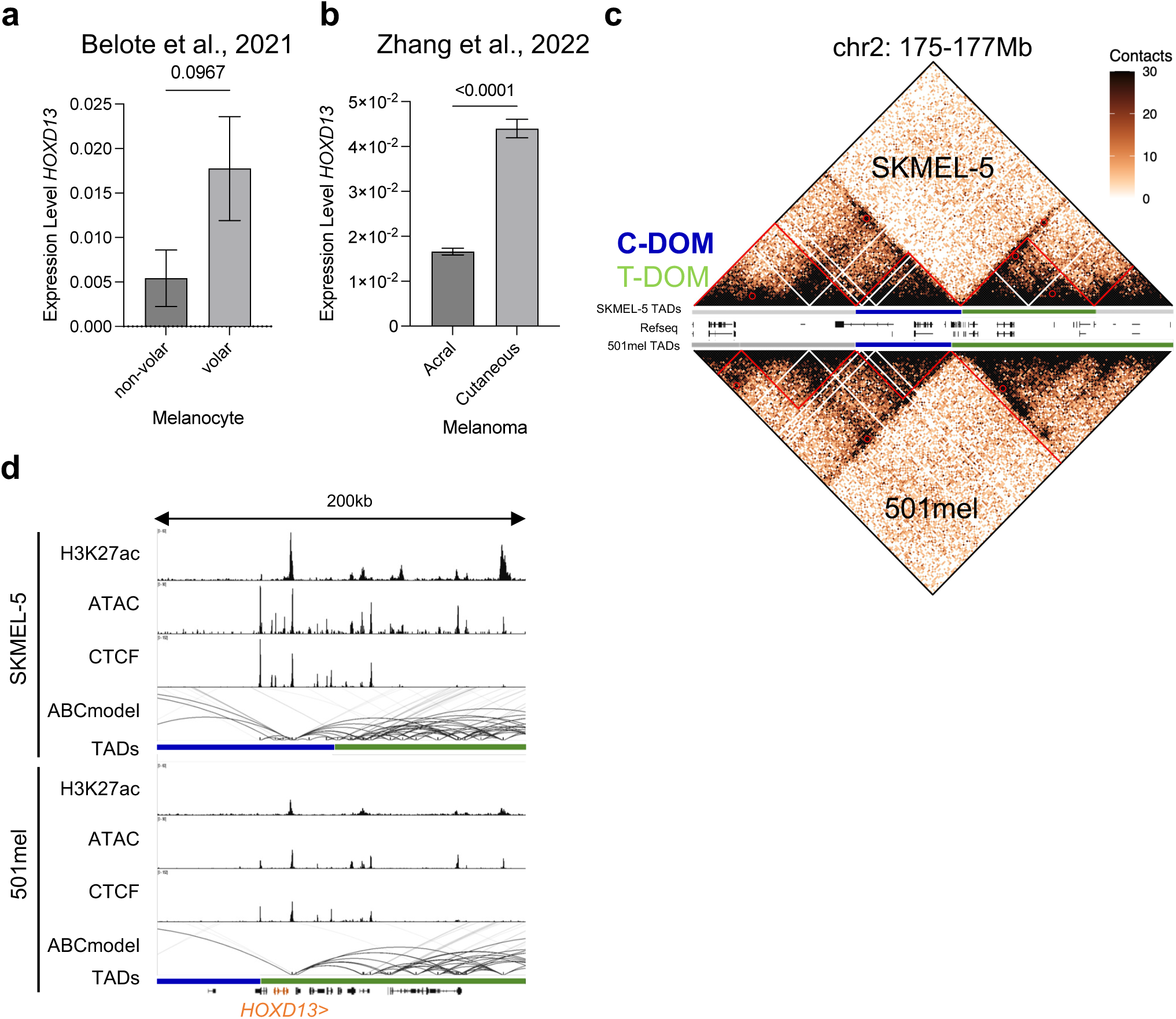
**a,** Violin plot showing *HOXD13* expression in human melanocytes by scRNAseq. No significant difference is observed between volar and non-volar melanocytes. Unpaired t test generated the p value. **b,** scRNAseq quantification of *HOXD13* expression between cutaneous and acral malignant melanoma cells demonstrate a significant upregulation of *HOXD13* in cutaneous vs acral. Unpaired t test was performed, and p value is shown. **c**, HiC pyramid plot showing the normalized contacts in a 2Mb window around *HOXD13* locus in SKMEL-5 (top) and 501mel (bottom). In red are marked the TADs. In the middle are shown the TADs coordinates and the gene bodies. **d**, IGV snapshot showing the H3K27ac, ATAC, CTCF, ABC model and TADs coordinates in a 200kb window around *HOXD13* locus. In blue and green are shown C-DOM and T-DOM respectively. In SKMEL-5 C-DOM is shifted to the right allowing the inclusion of HOXD13 to regulatory contacts. On the contrary, *HOXD13* gene is embedded in T-DOM which prevents upstream regulatory loops’ contacts and activation in 501mel.

